# Metabolomic, Lipidomic, and Enterohormone Changes in the Progression from MASLD to MASH

**DOI:** 10.1101/2025.09.16.676598

**Authors:** Jaclyn A. Rivas, Alexandria C. Murphy, Praveena Prasad, Siem S. Goitom, Aaron S. Romero, Crystal Madera Enriquez, Brianna B. Maes, Prithvi R. Akepati, Marcus A. Garcia, Fredine T. Lauer, Rama R. Gullapalli, Kristen M. Gonzales, Jessica M. Gross, Jing Pu, Shuguang Leng, Julie G. In, Melanie R. McReynolds, Eliseo F. Castillo

**Author notes:** Authors contributed equally to this paper (Jaclyn A. Rivas, Alexandria C. Murphy). Correspondence should be addressed to (E.F.C.) or (M.R.M.) 1 University of New Mexico Division of Gastroenterology and Hepatology Department of Internal Medicine University of New Mexico Health Sciences Center Albuquerque, NM 87131, USA.

## Abstract

**Background & Aims:** Metabolic Dysfunction-Associated Steatotic Liver Disease (MASLD) and Metabolic Dysfunction-Associated Steatohepatitis (MASH) represent progressive stages of liver disease, with distinct metabolic and cellular alterations. This study investigates the progression from MASLD to MASH through metabolomics, lipidomics, and assessment of hormones.

**Methods:** Male C57BL/6NTac mice were fed a high-fat diet for 16 weeks to induce MASLD and for 29 weeks to develop MASH. Aged-matched controls on a normal diet were used for comparison. Histology confirmed the progression of MASLD to MASH. We performed metabolomic and lipidomic profiling of liver, colon, and stool samples to identify metabolic and lipid alterations. Plasma enteroendocrine hormones and cytokines were quantified. Immunofluorescence was performed to assess enteroendocrine cells changes in the colon and the association of serotonin (5-HT) with fibronectin in the liver.

**Results:** Metabolomic and lipidomic analysis revealed significant alterations at different stages of the disease. Specifically, cholic acid was increased across the liver, colon, and stool in both MASLD and MASH mice compared to controls. Compared to the control group, MASLD mice exhibited an increase in enteroendocrine hormones, GLP-1, GIP, and PYY, whereas no changes were observed in MASH mice. Comparing MASLD to MASH livers, we found hepatic 5-HT levels were increased in MASH mice compared to MASLD mice. The MASH liver also exhibited a colocalization between fibronectin and 5-HT, suggesting a potential role of 5-HT in liver fibrosis.

**Conclusions:** Our study provides novel insights into the progressive metabolic and hormonal changes from MASLD to MASH. The increase in cholic acid and differential enteroendocrine hormone responses highlight the complex interactions between the gut and liver in metabolic liver diseases. These findings suggest that enteroendocrine hormones may play a role in the progression of MASLD to MASH as well as liver fibrosis, offering potential therapeutic avenues for targeting the gut-liver axis in metabolic liver diseases.

## Introduction

Metabolic Dysfunction-Associated Steatotic Liver Disease (MASLD) and its more advanced form, Metabolic Dysfunction-Associated Steatohepatitis (MASH), are emerging global health concerns driven by the obesity epidemic [1]. MASLD is characterized by hepatic fat accumulation, while MASH is marked by inflammation, hepatocyte injury, and fibrosis, potentially progressing to cirrhosis and hepatocellular carcinoma [2]. Despite the rising incidence of these diseases, the mechanisms driving the progression from simple steatosis (MASLD) to more severe steatohepatitis (MASH) remain poorly understood.

Enteroendocrine cells (EECs), which release hormones like glucagon-like peptide-1 (GLP-1) and glucose-dependent insulinotropic polypeptide (GIP) to regulate metabolism and liver function, yet their precise role in the progression from MASLD to MASH is still unclear [3–8]. Recent therapeutic advancements have focused on targeting metabolic regulation and liver inflammation, with GLP-1 receptor agonists, such as liraglutide and semaglutide, emerging as promising treatments. These therapies have demonstrated improved insulin sensitivity, reduce hepatic steatosis, and attenuate liver inflammation in both MASLD and MASH [9–11]. This success has translated clinically, as GLP-1 and GIP receptor dual agonists, like tirzepatide, have shown the potential to reduce hepatic fat accumulation and inflammation while improving insulin sensitivity [12, 13].

Other enteroendocrine hormones, such as serotonin (5-HT) produced by enterochromaffin cells (a subtype of EECs) [14, 15], have also emerged as an important signaling molecule in peripheral organs, including the gut and liver. Elevated 5-HT levels have been linked to liver fibrosis in chronic liver diseases, suggesting that 5-HT may contribute to the progression of MASH [16–18]. Despite these therapeutic advancements of these agents and the potential of some hormones to influence disease progression, the exact mechanisms particularly those involving enteroendocrine cell function and hormone release in fatty liver disease progression, remain incompletely understood.

The gut-liver axis, the bidirectional communication between the gut and liver, plays a critical role in fatty liver disease progression. This study aims to investigate the metabolic and hormonal changes that occur in the gut and liver during the progression from MASLD to MASH, with a particular focus on EECs. By employing metabolomic and lipidomic profiling, organoid and histological analysis, we found changes in hormones, lipids and metabolites that may drive the progression from MASLD to MASH.

## Materials and methods

Detailed ‘Materials and methods’ are provided in the supplementary information.

## Results

### Induction of MASLD and MASH with a modified amylin liver NASH diet

Diet-induced steatosis was established by feeding C57BL/6NTac male mice a modified amylin liver NASH diet (described in methods and herein referred to as high-fat diet, HFD) [19, 20], which substitutes palm oil for trans fats to mimic the absence of trans fat in the human diet. Starting at 6 weeks of age, mice were put on the HFD for 16 weeks to establish MASLD (HFD-16) and 29 weeks to establish MASH (HFD-29). Age-matched male mice (B6-16 or B6-29) were fed the NIH-31M chow diet to serve as a control group. As expected, mice on the HFD showed increased body weights (**Fig. 1A, F**) and liver weights (**Fig. 1B, G**) compared to the age-matched controls. Histological analysis of the liver from HFD-16 mice revealed steatosis (**Table S1** and **Fig. 1K, L**), consistent with the development of MASLD. Livers from HFD-29 mice showed more severe pathology, including a higher grade of steatosis, inflammation, hepatocellular hypertrophy, and fibrosis (**Table S1** and **Fig. 1M, N**), indicative of the progression to MASH. Inguinal fat pad weights were increased in HFD-16 compared to controls (**Fig. 1C**), although there were no significant weight differences in the inguinal fat pads when comparing HFD-29 and B6-29 (**Fig. 1H**). There were no observed differences in colon length between groups (**Fig. 1D, I**). Histological assessment also showed no intestinal inflammation in HFD-16 or HFD-29 mice compared to age-matched controls (**Table S2** and **Supplemental Fig. 1A-D**). Stool analysis for lipocalin-2 (LCN-2), a marker of intestinal inflammation [21, 22], showed no significant differences between HFD-16 and B6-16 mice (**Fig. 1E**), but an increase in HFD-29 mice (**Fig. 1J**). LCN-2 is a sensitive non-invasive marker of intestinal inflammation that increases even in the absence of clear histological signs of inflammation [23].

**Figure 1.**
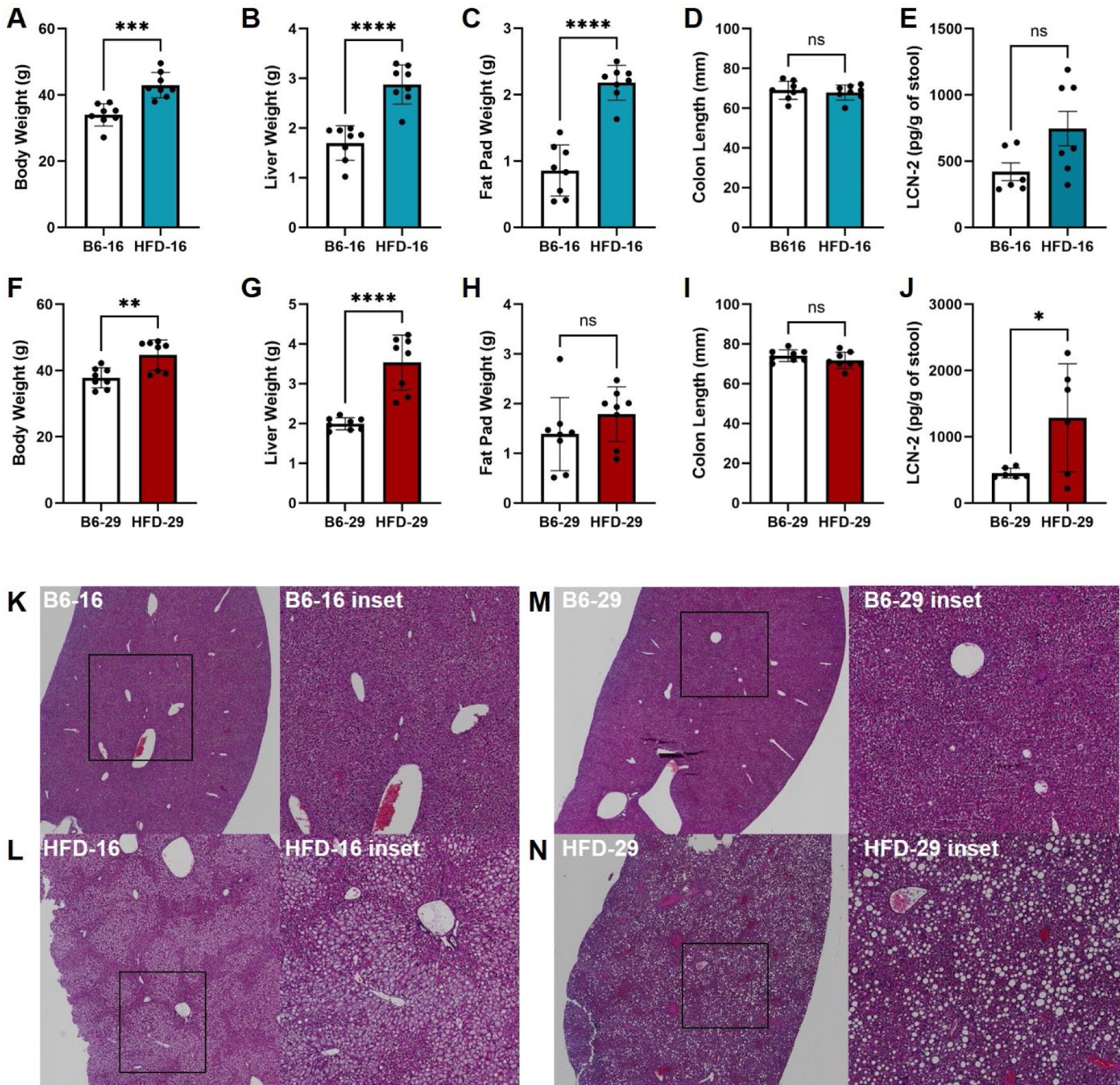
Gross assessment of diet-induced steatosis. C57BL/6NTac male mice were fed a modified Amylin liver NASH diet starting at 6 weeks of age to induce metabolic dysfunction-associated steatotic liver disease (MASLD) after 16 weeks (HFD-16) or metabolic dysfunction-associated steatohepatitis (MASH) after 29 weeks (HFD-29). Age-matched control mice (B6-16 and B6-29) were maintained on a standard NIH-31M chow diet. (**A-E)** Endpoint measurements in B6-16 vs HFD-16: (**A**) body weight, (**B**) liver weight, (**C**) fat pad weight, (**D**) colon length, and (**E**) fecal lipocalin-2 (LCN-2) levels. (**F-J)** Endpoint measurements in B6-29 vs HFD-29: (**F**) body weight, (**G**) liver weight, (**H**) fat pad weight, (**I**) colon length, and (**J**) fecal LCN-2 levels. (**K-N)** Representative H&E-stained liver sections from (**K**) B6-16, (**L**) HFD-16, (**M**) B6-29 and (**N**) HFD-29; histopathologic scoring summarized in Table S1. n = 8 mice per group. Data analyzed using Student’s t-test. *P < 0.05, **P < 0.005, ***P < 0.0005, \*\*\**P < 0.0001*.

### Metabolic profiles of the liver, colon, and stool from the MASLD mice

To map out metabolomics changes occurring between the gut and liver, untargeted metabolomics was performed on the liver and colon tissue and the stool collected from the colonic lumen across all groups. Key differences are summarized in **Table S3**. Metabolomics revealed significant alterations in the liver, colon, and stool of mice on a HFD compared to age-matched controls on a normal diet. For the HFD-16 group, several trends emerged in the data. In the colon (**Fig. 2A**), key metabolites like cholic acid, methylnicotinamide, and N-Me-2PY were significantly increased, while D-Phenyllactic acid, indole-3-ethanol, and other amino acid-related metabolites were significantly decreased. Volcano plots for the liver (**Fig. 2C**) show metabolites such as cholic acid, nicotinamide, and sucrose were significantly increased, while metabolites including fumarate, indole-3-ethanol, and retinal were notably decreased. In stool samples, metabolites such as cholic acid, cytidine, and 2-PY were significantly elevated, while kynurenic acid, melatonin, and sucrose were reduced (**Supplementary Fig. 3A**). Overall, the HFD-16 group exhibited notable disruptions in bile acid metabolism, amino acid homeostasis, and energy-related metabolites.

**Figure 2.**
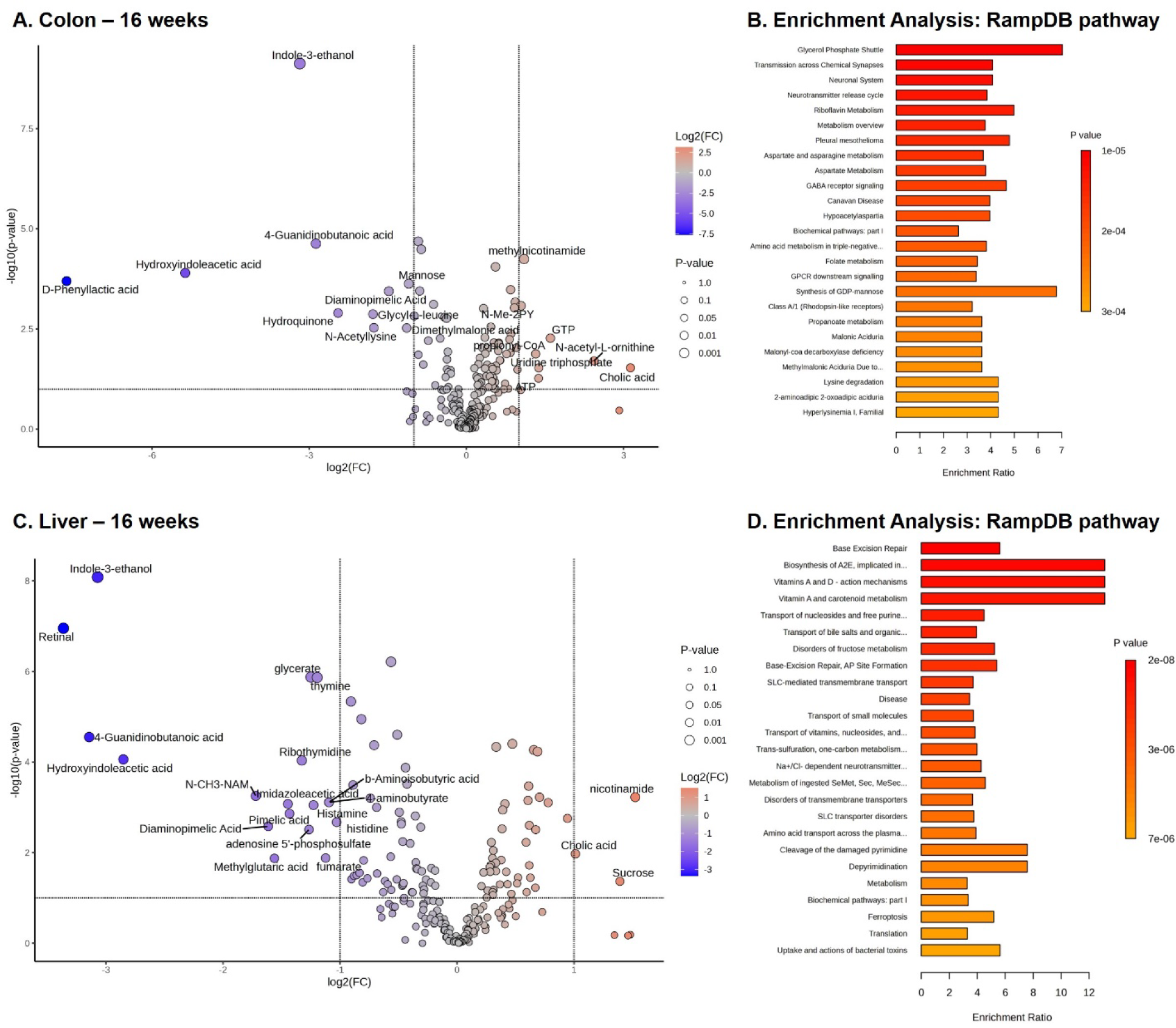
Metabolomic alterations in the colon and liver of HFD-fed mice at 16 weeks. (**A**, **C**) Volcano plots show significantly altered metabolites in the colon (**A**) and liver (**C**) of mice fed a high-fat diet (HFD-16) compared to age-matched controls on a normal chow diet. Log_2_ fold change (x-axis) is plotted against –log_10_ (p-value) (y-axis), with dot size indicating significance level and color indicating directionality of change. (**B**, **D**) Metabolite enrichment analysis using RaMP-DB highlights pathway-level changes in colon (**B**) and liver (**D**) metabolomes. Color denotes p-value, and bar length indicates enrichment ratio.

To further characterize the alterations in metabolites, pathway enrichment analysis was performed using the RaMP-DB (Relational database of Metabolomic Pathways) multi-sourced integrated database using multiple disease and pathway enrichment databases [24]. Metabolite enrichment analysis using RaMP-DB revealed significant pathway-level alterations in the colons of HFD-fed mice (**Fig. 2B**). The top enriched pathways included the glycerol phosphate shuttle, neurotransmitter-related signaling (e.g., GABA receptor signaling, transmission across chemical synapses, and the neuronal system), and amino acid metabolism (e.g., aspartate, folate, and lysine degradation) showing a HFD induces a broad metabolic reprogramming in colonic tissues, with potential impacts on host–microbiome interactions and epithelial signaling networks. To assess disease relevance, mouse colonic metabolites were matched to known human fecal or blood metabolite disease signatures, providing translational context by linking metabolic changes to gastrointestinal, hepatic, and systemic disease phenotypes. Analysis of our colon metabolites to the fecal signature revealed significant overlap with metabolites associated with colorectal cancer, Crohn’s disease, ulcerative colitis, and irritable bowel syndrome, as well as inflammatory conditions such as ankylosing spondylitis and rheumatoid arthritis (**Supplementary Fig. 2A**). Similarly, blood metabolite signature mapping revealed associations with cirrhosis, colorectal cancer, primary biliary cirrhosis, Alzheimer’s disease, and peroxisomal disorders (**Supplementary Fig. 2B).**

For the liver, pathway enrichment analysis using RaMP-DB revealed strong signatures of liver-specific metabolic processes, including bile acid metabolism, alcohol degradation, vitamin A metabolism, branched-chain and aromatic amino acid metabolism, fatty acid oxidation, and urea cycle pathways (**Fig. 2D**), all of which are central hepatic functions. Correspondingly, the fecal metabolite disease signature was enriched for obesity and several gut-related conditions: ulcerative colitis, Crohn’s disease and celiac disease, as well as inflammatory conditions such as ankylosing spondylitis and rheumatoid arthritis (**Supplementary Fig. 1C**). The blood metabolite disease signature further reinforced this association, with significant enrichment for obesity, adrenoleukodystrophy, peroxisomal biogenesis defect, and D-bifunctional protein deficiency (**Supplementary Fig. 1D**).

Metabolomic profiling of stool samples from mice with MASLD revealed extensive alterations in host and microbial metabolism, with pronounced enrichment of pathways linked to lipid signaling, neurotransmission, and inflammation (**Supplementary Fig. 3**). RaMP-DB pathway analysis identified strong enrichment for tryptophan metabolism and its catabolic pathways leading to NAD⁺ (nicotinamide adenine dinucleotide) biosynthesis, as well as the kynurenine pathway (**Supplementary Fig. 3B**). Additional pathways included glycosphingolipid metabolism, vitamin C metabolism, and eicosanoid synthesis pointing to alterations in lipid mediator production and oxidative stress responses. Notably, pathways involving G protein-coupled receptor (GPCR) signaling, neurotransmitter release, and gene expression regulation were also enriched, highlighting a potential gut-brain axis signaling. Disease signature analysis of fecal metabolites further supported our colon and liver findings, showing enrichment for gastrointestinal and inflammatory conditions, including ulcerative colitis, Crohn’s disease, and diverticular disease, alongside liver conditions like nonalcoholic fatty liver disease (**Supplementary Fig. 3C**). Complementary analysis of blood metabolite signatures revealed links to diabetes mellitus type 2 and infantile liver failure syndrome 2 as well as systemic metabolic disturbances, particularly in pathways associated with mitochondrial function and lipid oxidation, as evidenced by enrichment for disorders such as 3-hydroxy-3-methylglutaryl-CoA lyase deficiency and propionic academia (**Supplementary Fig. 3D**). Together, these data reveal that early-stage MASLD is characterized by coordinated metabolic reprogramming across the gut–liver axis, with disruptions in bile acid and amino acid metabolism, microbial signaling, and lipid homeostasis that align with human gastrointestinal, hepatic, and systemic disease signatures.

### Metabolic profiles of the liver, colon, and stool from the MASH mice

For the HFD-29 group, the metabolic changes became more pronounced, reflecting the progression from MASLD to MASH. In the colon, key metabolites including cholic acid, glycerate, and N-acetyl-L-proline were significantly elevated, while butyric acid, D-Phenyllactic acid, and hydroquinone were some of the metabolites reduced (**Fig. 3A**). While no intestinal inflammation was observed by histology (**Supplementary Fig. 1**), the metabolic alterations suggest potential shifts in gut microbial activity and metabolism in response to the prolonged high-fat diet exposure. In the liver, metabolites such as cholic acid, cyclic AMP, and O-phosphoethanolamine were significantly increased, while metabolites like 4-Guanidinobutanoic acid, indole-3-ethanol, and pimelic acid were significantly decreased (**Fig. 3C**). In stool samples, cholic acid, 2-PY, and leucine were significantly elevated, while metabolites like glucose, kynurenic acid, and melatonin were reduced (**Supplementary Fig. 5A**).

**Figure 3.**
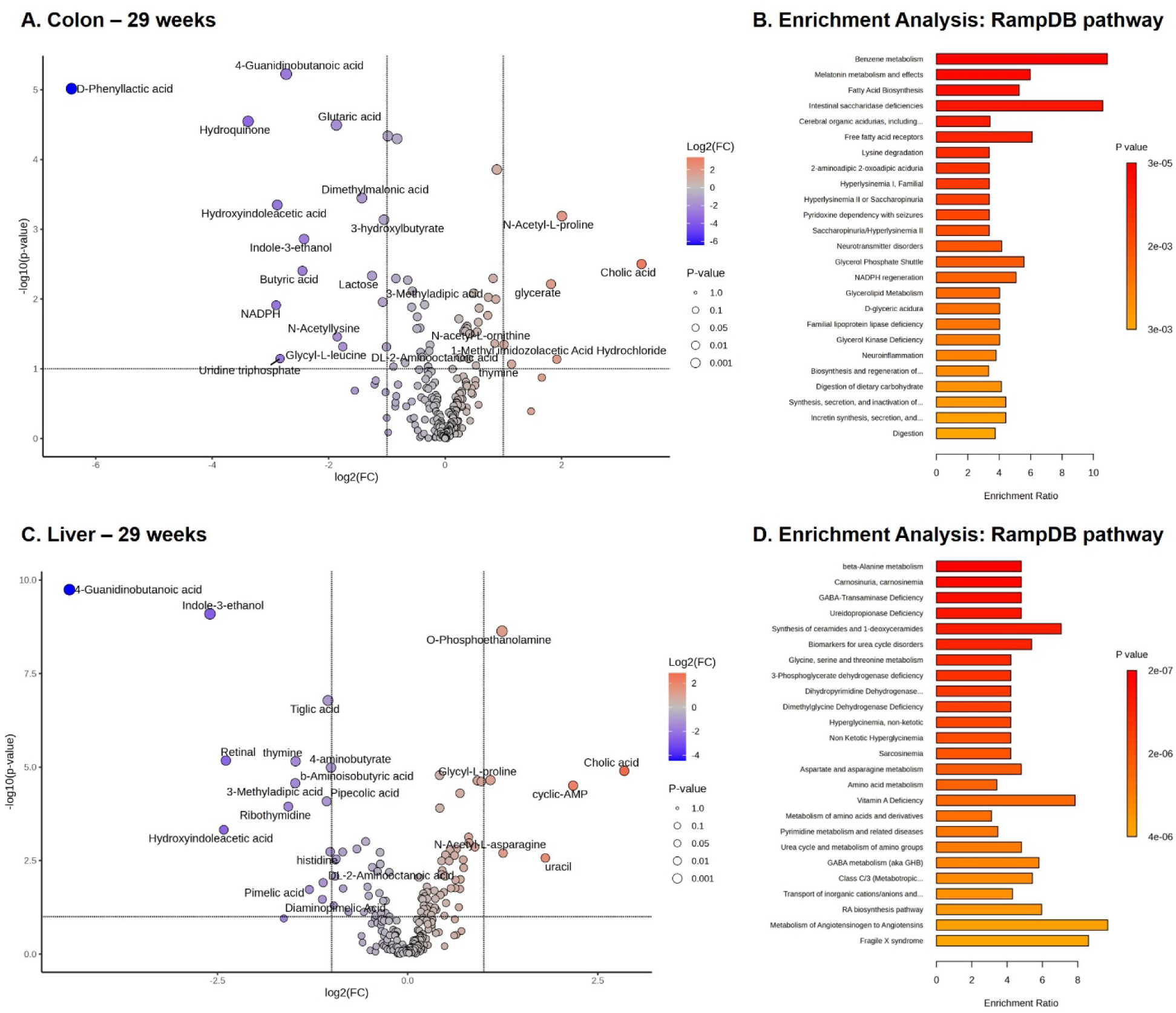
Metabolomic alterations in the colon and liver of HFD-fed mice at 29 weeks. (**A**, **C**) Volcano plots show significantly altered metabolites in the colon (**A**) and liver (**C**) of mice fed a high-fat diet (HFD-29) compared to age-matched controls on a normal chow diet. Log_2_ fold change (x-axis) is plotted against –log_10_ (p-value) (y-axis), with dot size indicating significance level and color indicating directionality of change. (**B**, **D**) Metabolite enrichment analysis using RaMP-DB highlights pathway-level changes in colon (**B**) and liver (**D**) metabolomes. Color denotes p-value, and bar length indicates enrichment ratio.

Pathway and disease enrichment analysis on metabolites derived from colon tissue using RaMP-DB, along with corresponding fecal and blood metabolite signatures were also assessed in MASH mice. RaMP-DB analysis revealed significant enrichment in pathways related to fatty acid biosynthesis, intestinal saccharidase deficiencies, benzene metabolism, and melatonin metabolism, indicating broad disruption of lipid processing, intestinal enzymatic activity, and host-microbe signaling pathways (**Fig. 3B**). Fecal metabolite signatures were enriched for disease phenotypes associated with intestinal and systemic inflammation, including *Clostridium difficile* infection, sepsis, colorectal cancer, and various enteric infections, highlighting the impact of MASH on gut microbiota-derived metabolism and intestinal barrier integrity (**Supplementary Fig. 4A**). Disease enrichment analysis of blood metabolites from MASH colons mice revealed rare peroxisomal and mitochondrial disorders, including D-bifunctional protein deficiency, peroxisomal biogenesis defects, and adrenoleukodystrophy (**Supplementary Fig. 4B**). These disorders are characterized by impaired fatty acid oxidation, disrupted lipid homeostasis, and bile acid metabolism abnormalities, consistent with the systemic metabolic stress observed in MASH.

In the liver, RaMP-DB pathway enrichment analysis highlighted significant perturbations in beta-alanine metabolism, carnosinuria/carnosinemia, and deficiencies related to GABA-transaminase and ureidopropionase (**Fig. 3D**). Additional enriched pathways included the synthesis of ceramides and 1-deoxyceramides, the urea cycle, and broader amino acid metabolic networks. Notably, pathways involved in the metabolism of angiotensinogen and the retinoic acid biosynthesis pathway were also significantly enriched, mirroring findings observed in the colon dataset and suggesting systemic metabolic involvement. The fecal metabolite disease signature from these liver-targeted metabolites was dominated by gastrointestinal and inflammatory diseases (**Supplementary Fig. 3C**). Top enrichments included Crohn’s disease, ulcerative colitis, and colorectal cancer, along with recurrent *Clostridium difficile* infection and irritable bowel syndrome. Interestingly, signatures for neurological and systemic conditions such as autism and myalgic encephalomyelitis were also present, further supporting the hypothesis of a gut-liver-brain axis playing a role in metabolic regulation and disease susceptibility. Mapping liver metabolites to the blood metabolite disease signature reflected a spectrum of liver-related and systemic conditions (**Supplementary Fig. 3D**). Cirrhosis was an enriched disease signature, accompanied by diseases such as cystic fibrosis, primary biliary cirrhosis, and uremia were prominent, indicating disruption of hepatobiliary and renal metabolic functions. Neurodegenerative and neuropsychiatric conditions, including Alzheimer’s disease and schizophrenia, were also enriched, suggesting broader systemic effects.

For the stool metabolite profile of MASH mice revealed a distinct disease and pathway enrichment signature indicative of systemic metabolic dysfunction and gut-associated disorders. Enrichment analysis using RaMP-DB showed strong associations with pathways involved in neurotransmitter metabolism, including 5-HT, tryptophan, and kynurenine metabolism, as well as melatonin metabolism and glycosphingolipid metabolism (**Supplementary Fig. 5B**). Feces disease signature analysis revealed significant enrichment in conditions related to gut inflammation and dysbiosis, including ulcerative colitis, Crohn’s disease, *Clostridium difficile* infection, and colorectal cancer (**Supplementary Fig. 5C**). Additionally, immune-related conditions such as sepsis and iron deficiency further underscore the inflammatory and immunometabolic disturbances linked with MASH. In comparison, the blood-derived metabolite signature highlighted metabolic disorders including NAD deficiency, infantile liver failure syndrome, and congenital adrenal hyperplasia, with strong enrichment in mitochondrial and peroxisomal dysfunction pathways (**Supplementary Fig. 5D**). Collectively, these data demonstrate that MASH is characterized by extensive metabolic dysregulation across the gut-liver axis, involving disrupted lipid and amino acid metabolism, neurotransmitter pathways, and systemic metabolic stress linked to mitochondrial and peroxisomal dysfunction, with metabolite signatures associated with intestinal barrier impairment, inflammation, and neuroimmune disorders.

### Tissue-Specific Lipid Remodeling and Functional Disruption in the Colon and Liver during Early Fatty Liver Disease

To investigate lipidomic changes associated with the HFD-induced MASLD (HFD-16), we performed untargeted lipidomics on the colon and liver. Compared to age-matched chow-fed B6-16, MASLD mice exhibited broad alterations in lipid class composition, with the liver showing a wider range of changes than the colon. In the colon, multiple lipid classes were elevated, including diacylglycerols (DG), lysophosphatidylcholines (LPC), lysophosphatidylethanolamines (LPE), phosphatidylinositols (PI), phosphatidylserines (PS), and hemibismonoacylglycerophosphates (HBMP). These classes support membrane curvature, vesicle trafficking, and signal transduction, suggesting early epithelial remodeling and altered host–microbe communication. In contrast, sulfonolipids (SL)-sphingolipid-like molecules involved in microbial signaling and epithelial homeostasis were markedly reduced, hinting at compromised gut barrier function or disrupted microbial-derived lipid signaling (**Fig. 4A, B**).

**Figure 4.**
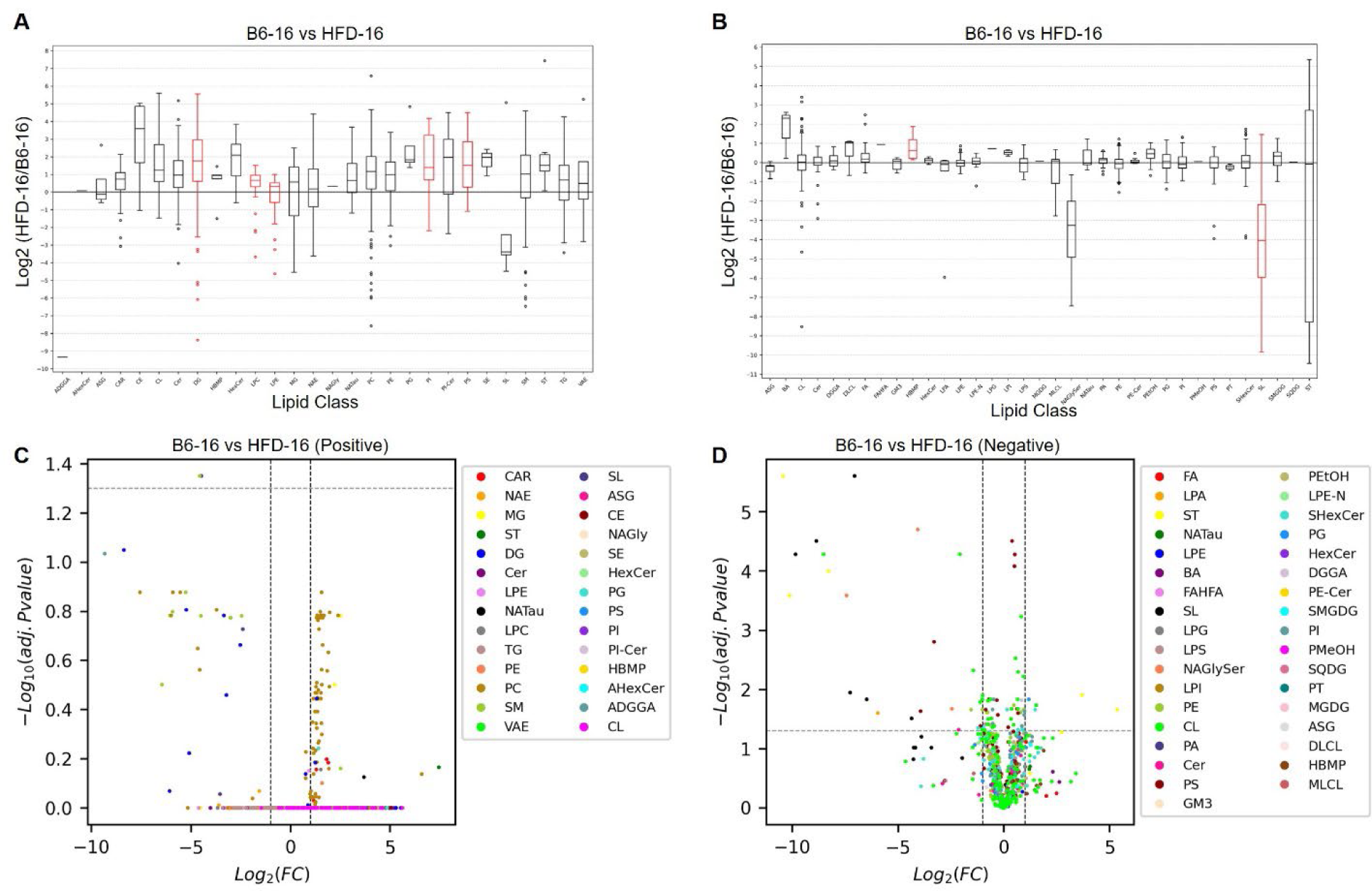
Lipidomic alterations in the colon of HFD-fed mice at 16 weeks. **(A)** The box plot shows the log2-fold change in lipid class enrichment in the colon of HFD-fed mice (HFD-16), with age-matched controls on normal chow diet as the reference (positive ionization mode). Red boxes are significant and black boxes are not significant. (**B**) The box plot shows the log2-fold change in lipid class enrichment in the colon of mice fed a high-fat diet (HFD-16), with age-matched controls on normal chow diet as the reference (negative ionization mode). Red boxes are significant and black boxes are not significant. (**C**) Volcano plots show significantly altered lipids in the colon of mice fed a high-fat diet (HFD-16) compared to age-matched controls on a normal chow diet. Log2 fold change (x-axis) is plotted against –log10 (p-value) (y-axis), with dot size indicating significance level and color indicating directionality of change (positive ionization mode). (**D**) Volcano plots show significantly altered lipids in the colon of mice fed a high-fat diet (HFD-16) compared to age-matched controls on a normal chow diet. Log2 fold change (x-axis) is plotted against –log10 (p-value) (y-axis), with dot size indicating significance level and color indicating directionality of change (negative ionization mode). Table S4 details lipid abbreviation.

The liver lipidome showed even more extensive remodeling. MASLD livers accumulated phosphatidylethanolamines (PE), acylcarnitines (CAR), N-acylethanolamines (NAE), LPC, phosphatidylcholines (PC), hexosylceramides (HexCer), DG, LPE, HBMP, phosphatidylglycerols (PG), sterols (ST), and free fatty acids (FA). These changes are consistent with mitochondrial stress (e.g., elevated CAR), dysregulated lipid turnover (e.g., LPC, DG, PC), and activation of inflammatory or fibrotic pathways. In particular, NAEs and HexCer have been implicated in endocannabinoid and ceramide-mediated stress responses, respectively, which may contribute to early steatotic injury. Similar to the colon, SL levels were reduced, alongside decreases in monoacylglycerols (MG) and N-acyl glycyl serines (NaGlyser) (**Fig. 5A, B**), indicating a disruption in lipid-mediated regulatory networks across tissues.

**Figure 5.**
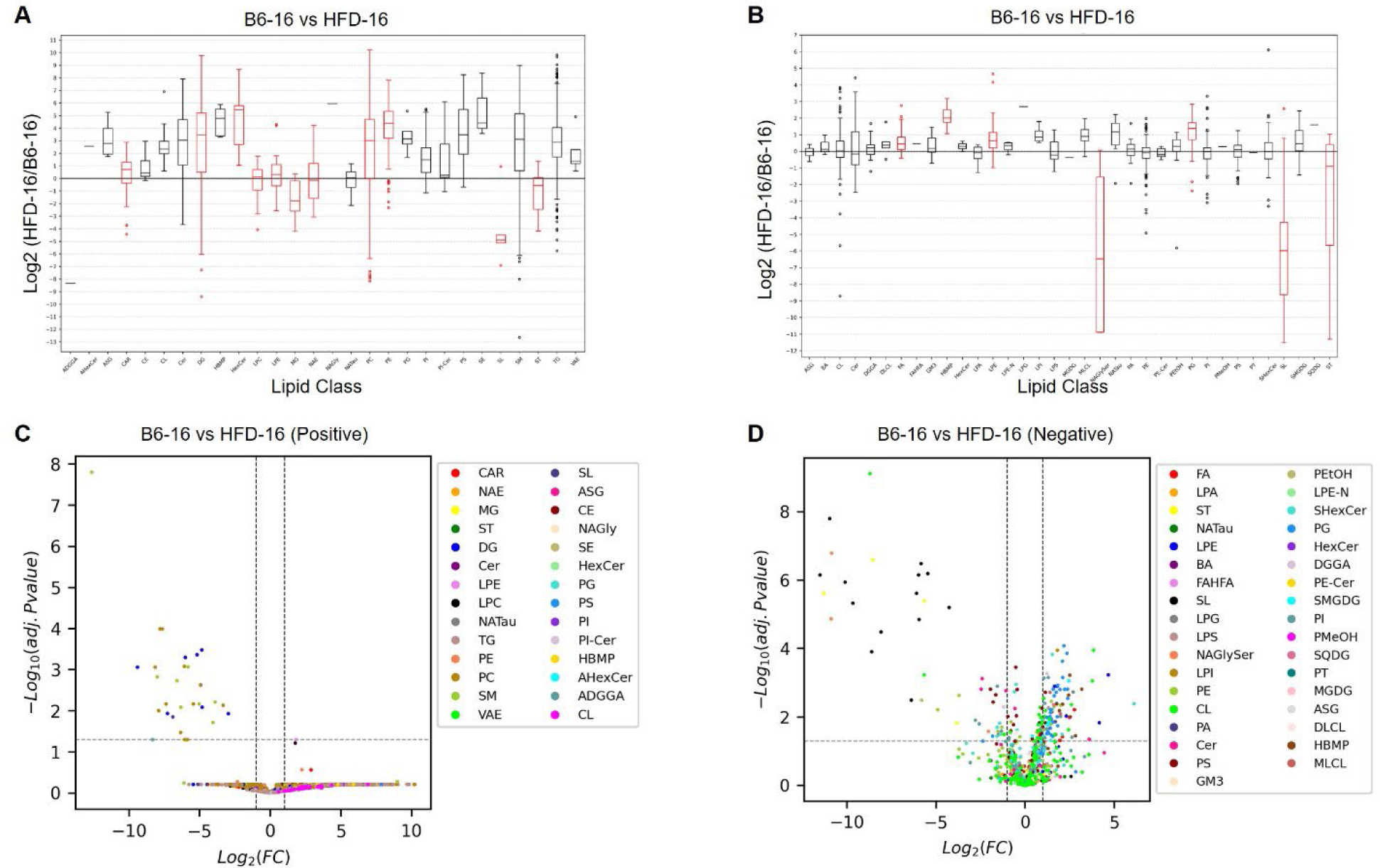
Lipidomic alterations in the liver of HFD-fed mice at 16 weeks. (**A**) The box plot shows the log2-fold change in lipid class enrichment in the liver of HFD-fed mice (HFD-16), with age-matched controls on normal chow diet as the reference (positive ionization mode). Red boxes are significant and black boxes are not significant. (**B**) The box plot shows the log2-fold change in lipid class enrichment in the liver of mice fed a high-fat diet (HFD-16), with age-matched controls on normal chow diet as the reference (negative ionization mode). Red boxes are significant and black boxes are not significant. (**C**) Volcano plots show significantly altered lipids in the liver of mice fed a high-fat diet (HFD-16) compared to age-matched controls on a normal chow diet. Log2 fold change (x-axis) is plotted against –log10 (p-value) (y-axis), with dot size indicating significance level and color indicating directionality of change (positive ionization mode). (**D**) Volcano plots show significantly altered lipids in the liver of mice fed a high-fat diet (HFD-16) compared to age-matched controls on a normal chow diet. Log2 fold change (x-axis) is plotted against –log10 (p-value) (y-axis), with dot size indicating significance level and color indicating directionality of change (negative ionization mode). Table S4 details lipid abbreviation.

Class-level shifts were corroborated by lipid-specific heatmaps and volcano plots for colon (**Supplementary Fig. 6C, D; Fig. 4C, D**) and liver (**Supplementary Fig. 7C, D; Fig. 5C, D**), which highlighted DG, PC, SL, and ST as consistently altered in both organs. Notably, MASLD livers also showed greater changes in lipid chain-length distribution (**Supplementary Fig. 6A, B vs. 7A, B**), potentially reflecting altered desaturation and elongation processes characteristic of metabolic stress and early fibrogenesis.

### Tissue-Specific Lipid Remodeling and Functional Disruption in the Colon and Liver during MASH

Untargeted lipidomic profiling of colon and liver tissues from MASH mice (HFD-29) revealed distinct, tissue-specific alterations in lipid class composition and chain length distribution compared to age-matched B6-29 mice. In the colon, levels of HexCer, LPE, PE, PC, and FA were significantly increased (**Fig. 6A, B**). These lipid classes participate in membrane remodeling, stress adaptation, and lipid-mediated signaling, suggesting compensatory epithelial remodeling in response to chronic injury. Conversely, sulfonolipids (SL), triacylglycerols (TG), and N-acyl glycyl serines (NaGlySer) lipids involved in microbial interactions, energy storage, and intercellular communication were markedly decreased.

**Figure 6.**
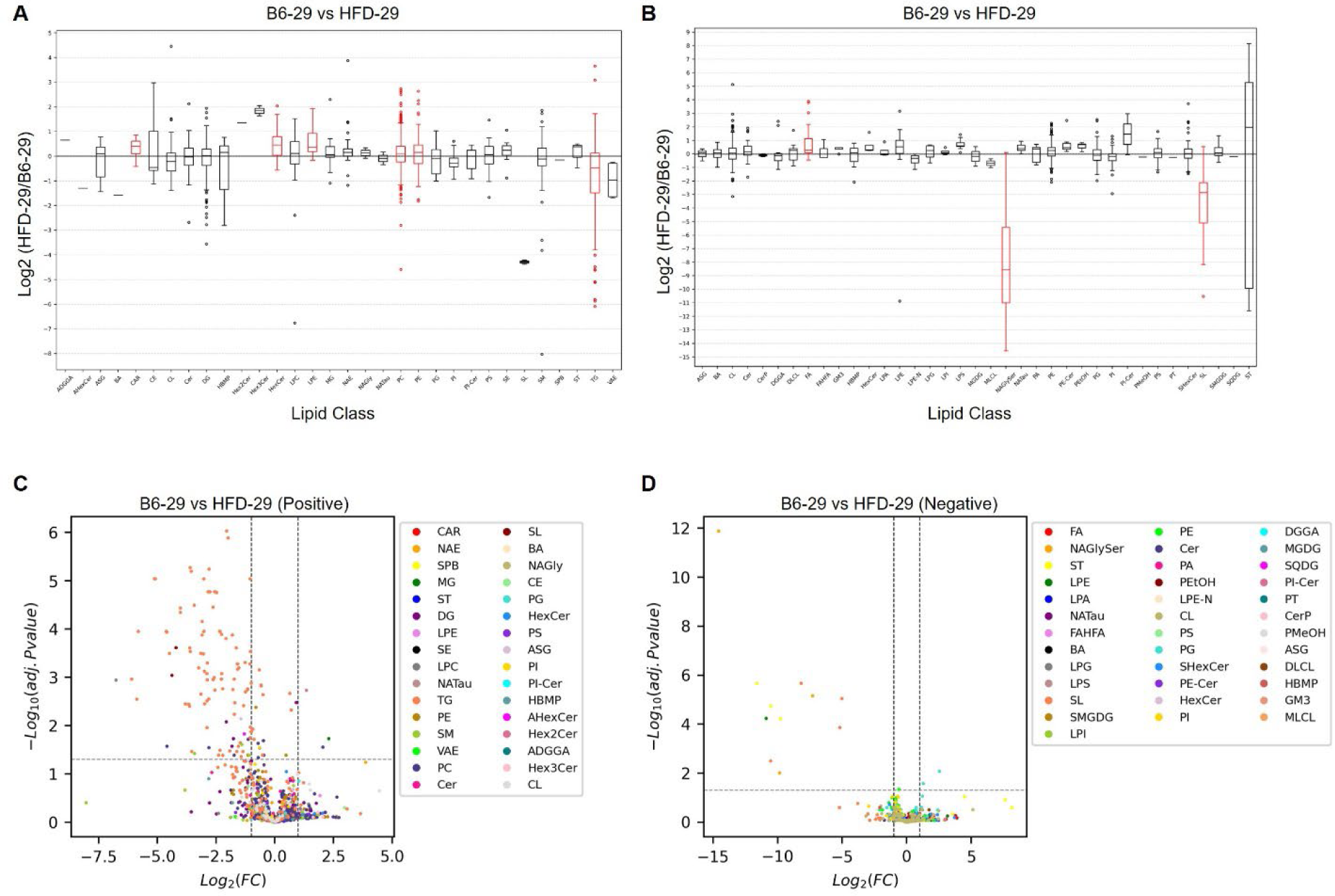
Lipidomic alterations in the colon of HFD-fed mice at 29 weeks. (**A**) The box plot shows the log2-fold change in lipid class enrichment in the colon of HFD-fed mice (HFD-29), with age-matched controls on normal chow diet as the reference (positive ionization mode). Red boxes are significant and black boxes are not significant. (**B**) The box plot shows the log2-fold change in lipid class enrichment in the colon of mice fed a high-fat diet (HFD-29), with age-matched controls on normal chow diet as the reference (negative ionization mode). Red boxes are significant and black boxes are not significant. (**C**) Volcano plots show significantly altered lipids in the colon of mice fed a high-fat diet (HFD-29) compared to age-matched controls on a normal chow diet. Log2 fold change (x-axis) is plotted against –log10 (p-value) (y-axis), with dot size indicating significance level and color indicating directionality of change (positive ionization mode). (**D**) Volcano plots show significantly altered lipids in the colon of mice fed a high-fat diet (HFD-29) compared to age-matched controls on a normal chow diet. Log2 fold change (x-axis) is plotted against –log10 (p-value) (y-axis), with dot size indicating significance level and color indicating directionality of change (negative ionization mode). Table S4 details lipid abbreviation.

In the liver, MASH was associated with elevated ceramides (CE), monoacylglycerols (MG), N-acylethanolamines (NAE), HBMP, PI, and PG (**Fig. 7A, B**). These changes implicate activation of pro-inflammatory and endocannabinoid signaling, as well as altered membrane trafficking. Acylcarnitines (CAR), markers of mitochondrial dysfunction and impaired β-oxidation, were increased in both colon and liver, consistent with systemic metabolic stress.

**Figure 7.**
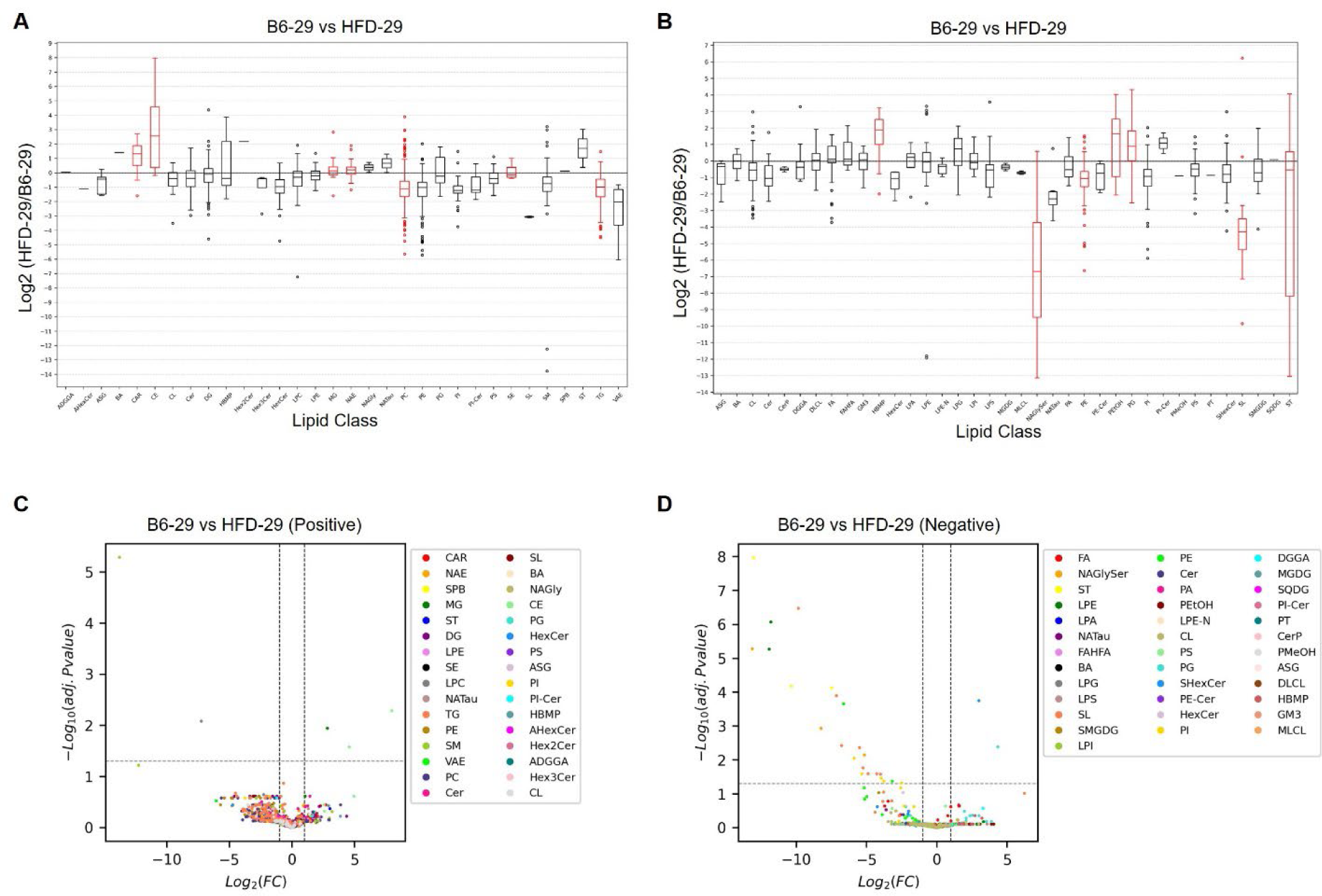
Lipidomic alterations in the liver of HFD-fed mice at 29 weeks. (**A**) The box plot shows the log2-fold change in lipid class enrichment in the liver of HFD-fed mice (HFD-29), with age-matched controls on normal chow diet as the reference (positive ionization mode). Red boxes are significant and black boxes are not significant. (**B**) The box plot shows the log2-fold change in lipid class enrichment in the liver of mice fed a high-fat diet (HFD-29), with age-matched controls on normal chow diet as the reference (negative ionization mode). Red boxes are significant and black boxes are not significant. (**C**) Volcano plots show significantly altered lipids in the liver of mice fed a high-fat diet (HFD-29) compared to age-matched controls on a normal chow diet. Log2 fold change (x-axis) is plotted against –log10 (p-value) (y-axis), with dot size indicating significance level and color indicating directionality of change (positive ionization mode). (**D**) Volcano plots show significantly altered lipids in the liver of mice fed a high-fat diet (HFD-29) compared to age-matched controls on a normal chow diet. Log2 fold change (x-axis) is plotted against –log10 (p-value) (y-axis), with dot size indicating significance level and color indicating directionality of change (negative ionization mode). Table S4 details lipid abbreviation.

Volcano plot analyses supported these trends, showing decreased TG, SL, and PI in the colon (**Fig. 6C, D**) and SL and PI in the liver (**Fig. 7C, D**). Liver-specific reductions in PE and sterols (ST) suggest perturbed membrane composition and cholesterol metabolism. Notably, in contrast to MASLD, where lipid remodeling was more pronounced in the liver, MASH mice displayed greater shifts in lipid chain length in the colon (**Supplementary Fig. 8A, B**) whereas the liver exhibited fewer chain length alterations (**Supplementary Fig. 9A, B**). Together, these results indicate that progression from MASLD to MASH involves a redistribution of lipidomic stress from the liver to the colon, reflecting dynamic reprogramming of lipid signaling networks, mitochondrial function, and potentially gut-liver communication during advanced disease.

### Alterations in circulating enteroendocrine cell hormones

Metabolomic analysis revealed increased levels of cholic acid across all tissue types in both MASLD and MASH samples (**Tables S3**). Multiple studies have demonstrated that bile acids, including cholic acid, can activate EECs via takeda G protein-coupled receptor 5 (TGR5), a GPCR [25–29]. Supporting this, our pathway analysis indicated upregulation of signaling pathways associated with EEC function, including “Incretin synthesis, secretion, and inactivation”, “Synthesis, secretion, and inactivation of GLP-1” as well as “GPCR downstream signaling” and “GPCR ligand binding” in both colon and stool samples.

Based on these findings, we next examined circulating levels of enteroendocrine hormones, metabolic regulators as well as pro-inflammatory cytokines across our study groups. Plasma concentrations of GIP, GLP-1, peptide tyrosine tyrosine (PYY), ghrelin, secretin, 5-HT, insulin, amylin, pancreatic peptide (PP), C-peptide, leptin, and resistin were measured in HFD-16 and HFD-29 mice and compared to age-matched controls. Additionally, we quantified levels of the inflammatory cytokines tumor necrosis factor (TNF), interleukin-6 (IL-6), and monocyte chemoattractant protein 1 (MCP-1). In HFD-16 mice, we observed significant increases in GLP-1 (**Fig. 8 A**), GIP (**Fig. 8 B**), and PYY (**Fig. 8 C**) compared to age-matched controls with no change in 5-HT (**Fig. 8 D**), ghrelin (**Supplementary** Fig. 10 **A**) or secretin (**Supplementary** Fig. 10 **C**). In contrast, HFD-29 mice showed no significant difference in GLP-1 (**Fig. 8 E**), GIP (**Fig. 8 F**), PYY (**Fig. 8 G**), 5-HT (**Fig. 8 H**), ghrelin (**Supplementary Fig. 10 B**) or secretin (**Supplementary Fig. 10 D**) compared to age-matched controls. Notably, colonic cholic acid levels were positively correlated with GLP-1 (r = 0.6936; p = 0.0041) (**Fig. 8I**) and GIP levels (r = 0.5868; p = 0.0214) (**Fig. 8J**) in HFD-16 mice; however, no such correlation was observed in the HFD-29 group (data not shown).

**Figure 8.**
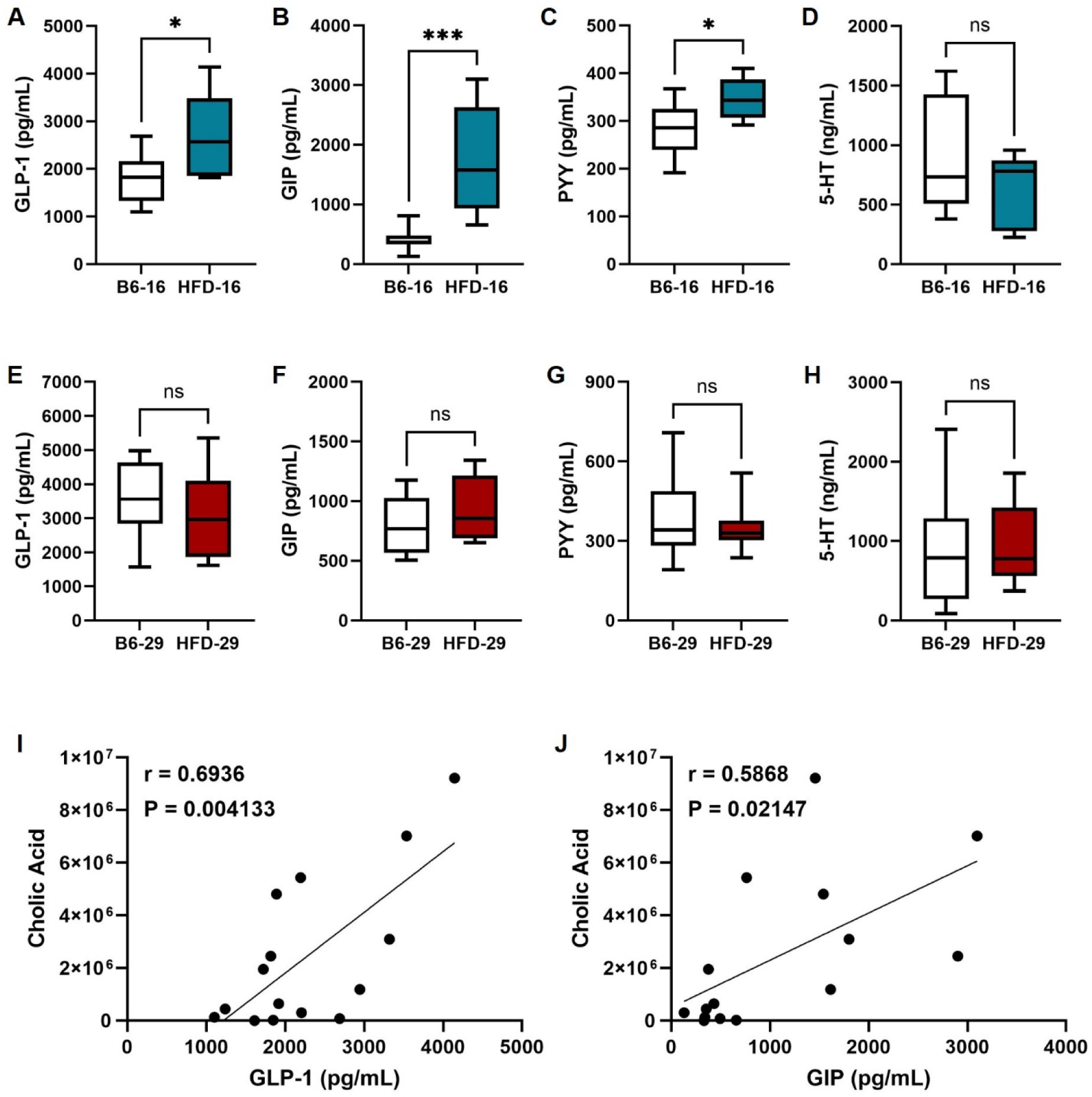
Diet-induced changes in enteroendocrine hormones and their correlation with cholic acid levels. (**A**-**H**) Circulating levels of enteroendocrine hormones were measured in plasma from mice fed a HFD for 16 weeks (HFD-16) or 29 weeks (HFD-29) and compared to age-matched controls on a normal diet (B6-16 or B6-29). (**A**-**D**) B6-16 versus HFD-16 hormones (**A**) GLP-1, (**B**) GIP, (**C**) PYY, and (**D**) 5-HT. (**E**-**H**) B6-29 versus HFD-29 hormones (**E**) GLP-1, (**F**) GIP, (**G**) PYY, and (**H**) 5-HT. (**I**, **J**) Colonic cholic acid levels positively correlated with (**I**) GLP-1 (r = 0.6936; P = 0.004133) and (**J**) GIP (r = 0.5868; P = 0.02147) levels in HFD-16 mice. *P < 0.05, ***P < 0.0005, NS = not significant.

We also found that HFD-16 mice had elevated plasma levels of insulin (**Supplementary Fig. 10 E**), amylin (**Supplementary Fig. 10 G**), leptin (**Supplementary Fig. 10 M**), resistin (**Supplementary** Fig. 10 **O**), and TNF (**Supplementary Fig. 10 Q**) compared to age-matched controls. Conversely, no differences in PP (**Supplementary Fig. 10 J**), C-peptide (**Supplementary Fig. 10 K**), IL-6 (**Supplementary Fig. 10 S**), or MCP-1 (**Supplementary Fig. 10 U**) were observed between groups. In HFD-29 mice, only TNF (**Supplementary Fig. 10 R**) and IL-6 (**Supplementary** Fig. 10 **T**) were significantly increased compared to age-matched controls with no differences in insulin, amylin, PP, C-peptide, leptin, resistin or MCP-1 (**Supplementary Fig. 10 F, H, J, L, N, P, and V**). In summary, circulating levels of GLP-1, GIP, and PYY were significantly elevated in MASLD (HFD-16) mice and positively correlated with colonic cholic acid levels. These hormone levels were unchanged in MASH (HFD-29) mice, despite persistently elevated cholic acid, suggesting stage-specific alterations in EEC responsiveness. Furthermore, MASH mice exhibited increased TNF and IL-6 levels, indicative of a shift toward chronic inflammation with dampened incretin and metabolic hormone responses in later disease stages.

To determine whether these stage-specific differences in EEC abundance reflected intrinsic changes in epithelial differentiation potential, we next used organoid cultures derived from colonic crypts of each group. We collected colonic crypts from all groups to generate organoids, which were initially maintained in the intestinal stem cell state and then induced to differentiate by withdrawing WNT3A and R-spondin. Successful differentiation was confirmed by the expected reduction in *Lgr5* expression, a widely used stem cell marker (**Supplementary Fig. 11A, B, F, G**). We next examined the expression of genes associated with the EEC lineage. For the 16 week groups, we observed no significant differences in *Neurog3*, *Chga*, or *Gcg* expression between organoids derived from B6-16 and HFD-16 mice (**Supplementary Fig. 11C–E**), indicating preserved EEC differentiation capacity at this stage. In contrast, at 29 weeks, organoids from HFD-fed mice showed a significant reduction in *Neurog3* and *Chga* expression compared to B6-29 controls, while *Gcg* expression remained unchanged (**Supplementary Fig. 11H–J**). These findings suggest that prolonged HFD exposure impairs EEC lineage specification and maturation, which may underlie the observed blunting of hormone responses *in vivo* during MASH progression. To further assess local EEC changes, we performed immunofluorescence staining for chromogranin A (CHGA), a general marker of EECs, in colonic tissue. HFD-16 mice exhibited an increase in CHGA⁺ cells within the colonic epithelium compared to age-matched controls, consistent with elevated circulating GLP-1 and PYY levels. In contrast, CHGA⁺ cell abundance in HFD-29 mice was similar to that of their age-matched controls, aligning with the lack of hormone elevation at this later disease stage (**Supplementary Fig. 11K–N**).

### Metabolic and 5-HT alterations in the liver between MASLD and MASH

We compared metabolite differences between MASLD and MASH livers. MASH livers were found to have a significant increase in cholic acid, cyclic-AMP, imidazoleacetic acid, linoleic acid, malate, 5’-Methylthioadenosine, serotonin, taurodeoxycholic acid, and uracil (**Fig. 9A**). Interestingly, linoleic acid [30–32], 5’-Methylthioadenosine [33, 34], and taurodeoxycholic acid [35, 36] have all been linked to enhanced liver disease and inflammation. Given the known role of enterochromaffin cell-derived 5-HT in gut-liver signaling and fibrogenesis, we next compared 5-HT levels between HFD-16 (MASLD) and HFD-29 (MASH) mice. Plasma 5-HT levels were higher in HFD-29 mice compared to HFD-16 mice, although this trend did not reach statistical significance (**Fig. 9B**). When comparing 5-HT concentrations across compartments, hepatic levels in HFD-29 mice were significantly elevated compared to HFD-16 (**Fig. 9C**). Metabolomic profiling confirmed a significant increase in hepatic 5-HT levels in HFD-29 mice relative to HFD-16 mice (**Fig. 9A**), a difference that was not observed in the colon (data not shown), suggesting local 5-HT accumulation or impaired hepatic clearance in the context of MASH.

**Figure 9.**
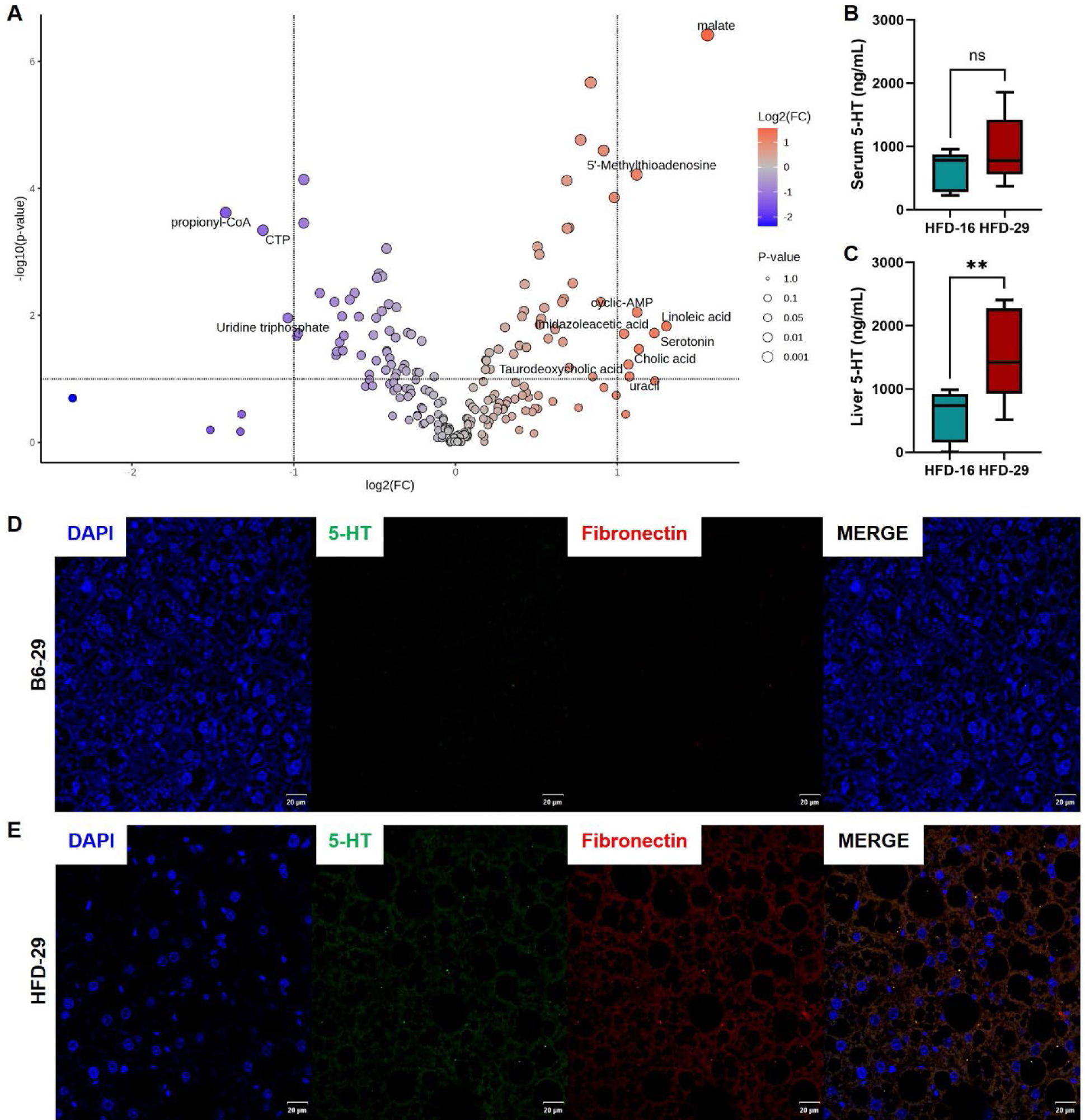
Hepatic 5-HT and Fibronectin distribution in MASH mice. (**A**, **B**) 5-HT levels as detected by ELISA were measured in (**A**) plasma and (**B**) liver tissues from HFD-16 (MASLD) and HFD-29 (MASH) mice. n = 8 mice per group. Data analyzed using Student’s t-test. **P < 0.005, NS = not significant. (**C**) Volcano plots show significantly altered hepatic metabolites in the HFD-29 compared to HFD-16 including 5-HT. Log_2_ fold change (x-axis) is plotted against –log_10_ (p-value) (y-axis), with dot size indicating significance level and color indicating directionality of change. (**D**, **E**) Representative immunofluorescence images of DAPI, 5-HT, and fibronectin staining of liver sections in (**D**) control B6-29 and (**E**) HFD-29 livers.

To examine whether this increase in hepatic 5-HT was associated with fibrotic remodeling, we performed immunofluorescence staining for fibronectin, a key extracellular matrix protein and marker of liver fibrosis. HFD-29 livers displayed marked fibronectin deposition (**Fig. 9E**), which was not detected in B6-29 controls (**Fig. 9D**). Importantly, 5-HT staining in HFD-29 livers showed partial co-localization with fibronectin-positive regions, suggesting that 5-HT accumulates within or contributes to fibrotic microenvironments (**Fig. 9E**). These findings raise the possibility that chronic HFD exposure leads to a shift in EEC-derived 5-HT distribution, promoting its hepatic retention and potential involvement in fibrosis during the transition from MASLD to MASH.

## Discussion

In this study, we leveraged a well-established dietary model to investigate how chronic HFD [19, 20] exposure drives progression from MASLD and MASH, with a specific focus on the gut-liver axis. Using integrated metabolomic, lipidomic, hormonal, and histological approaches, we demonstrate that this transition is marked by dynamic, stage-specific changes in EEC activity, metabolic hormone output, and 5-HT distribution. During early disease (MASLD), HFD-fed mice exhibited elevated levels of GLP-1, GIP, and PYY in the circulation, which correlated with increased cholic acid in the colon and liver and increased CHGA⁺ cell abundance in the colonic epithelium. Conversely, with prolonged HFD feeding and progression to MASH, these hormone elevations were lost despite persistent cholic acid elevation, suggesting a blunting or silencing of EEC responsiveness. This was supported by reduced expression of EEC lineage markers (*Neurog3*, *Chga*) in colonic organoids derived from HFD-29 mice. Concomitantly, hepatic 5-HT levels were significantly increased in MASH, with partial co-localization to fibronectin-rich regions, implicating 5-HT in fibrotic remodeling. This aligns with previous studies showing serotonin promotes hepatic stellate cell activation via the 5-HT2B receptor, stimulating TGF-β and collagen production [37–39]. These findings identify a temporal reprogramming of gut endocrine signaling across MASLD-to-MASH progression and highlight potential mechanistic links between EEC dysfunction, bile acid signaling, and hepatic fibrosis.

The observed changes in hormone output have important implications for systemic metabolic regulation. GIP and GLP-1 act as incretins that enhance insulin secretion following nutrient intake [40], while PYY contributes to appetite suppression and reduced gastrointestinal motility [41]. Insulin and amylin maintain glucose homeostasis [42]; leptin regulates energy balance and satiety [43]; and resistin has been linked to inflammation and insulin resistance [44, 45]. Observations in zebrafish show microbiota-dependent EEC silencing and highlight the dynamic plasticity of the gut endocrine system in response to chronic nutritional stress [46]. Our data reveal a time-dependent adaptation of enteroendocrine signaling to a HFD. At 16 weeks, HFD-fed mice exhibit elevated circulating levels of GLP-1, GIP, and PYY, consistent with EEC hyperactivation. Supporting this, organoids derived from HFD-16 and control mice showed no significant differences in the expression of EEC lineage genes (*Neurog3*, *Chga*, *Gcg*), suggesting intact differentiation capacity. However, by 29 weeks, hormone levels are comparable to age-matched controls, indicative of EEC desensitization or functional remodeling. In support of this, HFD-29-derived organoids show significant reductions in *Neurog3* and *Chga* expression, pointing to impaired EEC lineage commitment and maturation. Interestingly, 5-HT levels increase significantly in the liver at this stage, implicating a potential shift in EEC subtype activity, favoring enterochromaffin cell output or microbiota-mediated stimulation of 5-HT biosynthesis.

The results from the metabolomics analysis of liver, colon, and stool tissues highlight significant metabolic disturbances in mice on a HFD, with increasing severity from MASLD to MASH. The key trend of cholic acid accumulation across all tissues suggests an impairment in bile acid metabolism, potentially pointing to disrupted enterohepatic circulation and altered bile acid synthesis. Bile acids, which are synthesized in the liver and released into the small intestine, are crucial regulators of metabolic processes such as lipid metabolism, glucose homeostasis, and liver injury. Dysregulation of bile acid metabolism, both in the liver and the gut, has been implicated in the pathogenesis of MASLD and MASH [47, 48]. Altered bile acid profiles, including increased levels of cholic acid, have been associated with hepatic steatosis, inflammation, and fibrosis in both animal models [49–51] and human patients [52]. Furthermore, bile acids modulate the gut microbiota, influencing liver inflammation and fibrosis, suggesting a complex interaction between bile acid signaling and the gut-liver axis in disease progression. We hypothesize that bile acids may interact with EECs in the gut, stimulating the release of hormones such as GLP-1, PYY, and GIP that support lipid metabolism and hepatic function during early disease. Over time, however, persistent cholic acid elevation may contribute to EEC dysfunction or desensitization, disrupting gut hormone signaling and weakening the protective gut-liver feedback loop. This could facilitate progression from steatosis to steatohepatitis and systemic metabolic derangement. Additionally, both HFD groups showed downregulation of various amino acid metabolites and energy-related metabolites, with further dysregulation observed in the MASH group, indicating a progression toward more severe liver disease. The data also suggest that prolonged HFD exposure leads to gut microbiota changes, as evidenced by altered stool metabolite profiles. Overall, these findings provide novel insights into the metabolic shifts occurring in the liver, colon, and stool, and underscore the complex relationship between diet, metabolic dysregulation, and the progression of simple steatosis to steatohepatitis. Further investigation into the specific roles of these metabolites in disease progression could provide valuable targets for therapeutic intervention. While GLP-1 receptor agonists and dual GIP/GLP-1 agonists like tirzepatide have shown promise in improving metabolic parameters and reducing hepatic steatosis in preclinical and early clinical studies, their long-term efficacy in treating MASLD and MASH remains under active investigation. Clinical experience with these agents, particularly in liver disease, is still evolving, and questions remain regarding the magnitude and durability of their therapeutic effects on hepatic inflammation and fibrosis.

Lipidomic analysis of the liver and colon revealed dynamic, tissue-specific remodeling of lipid classes that parallels the progression from MASLD to MASH in HFD-fed mice. In the MASLD stage, the liver displayed a robust upregulation of free fatty acids (FAs), an early and well-documented hallmark of hepatic steatosis and lipid overload in MASLD pathogenesis[53]. As key precursors for complex lipid biosynthesis, the elevation of FAs is consistent with the metabolic shift toward hepatic lipid accumulation. Accompanying this, both the liver and colon showed increased levels of diacylglycerols (DG), a lipid class directly linked to hepatic insulin resistance and metabolic dysfunction[54], thereby reinforcing the connection between lipid dysregulation and disease progression. Additionally, MASLD tissues exhibited higher levels of lysophosphatidylcholine (LPC) and lysophosphatidylethanolamine (LPE), bioactive lipid classes known to induce lipotoxicity in hepatocytes and contribute to inflammatory signaling cascades [55]. Together, the upregulation of FAs, DG, LPC, and LPE in MASLD suggests a coordinated lipotoxic program that promotes metabolic stress and may predispose tissues to further injury and progression toward MASH.

In contrast, the MASH phenotype was marked by a distinct shift in lipid signatures. Notably, acylcarnitines (CAR) were the only lipid class consistently increased in both liver and colon of MASH mice. As intermediates of mitochondrial fatty acid oxidation, elevated CAR levels indicate mitochondrial overload or incomplete β-oxidation, which is a hallmark of advanced liver pathology. Furthermore, CAR accumulation has been associated with hepatocellular carcinoma (HCC), suggesting that their elevation in MASH may reflect a metabolic trajectory toward oncogenic transformation[56]. Thus, CAR species may serve as early biomarkers for MASH severity and its potential progression to liver cancer.

Additional patterns of lipid remodeling further distinguish MASLD from MASH. Phosphatidylethanolamines (PE), essential components of membrane structure and lipid bilayer integrity, were elevated in MASLD liver but significantly reduced in MASH liver, mirroring previous observations that PE depletion correlates with disease advancement [57]. A consistent downregulation of sulfonolipids (SL) in both liver and colon across disease stages provides further mechanistic insight into gut-liver communication. As microbiota-derived lipids structurally analogous to sphingolipids, SLs are emerging mediators of immune homeostasis and epithelial integrity [58]. Their depletion supports the hypothesis that microbial lipid signaling is disrupted in the context of MASLD/MASH, contributing to immune dysregulation and disease progression.

These lipid alterations may also influence bile acid composition and enteroendocrine cell function, potentially linking shifts in membrane remodeling and mitochondrial metabolism to changes in gut-liver hormonal and metabolic signaling. Together, these findings highlight that MASLD and MASH are characterized by distinct lipidomic signatures reflecting early lipotoxic stress (FAs, DG, LPC/LPE), mitochondrial dysfunction (CAR), and disrupted host–microbial signaling (SL). These tissue-specific lipid alterations offer mechanistic insight into the gut–liver axis in metabolic liver disease and may serve as candidate biomarkers for disease stage and therapeutic targeting.

### Limitation of Study

This study has several limitations. While we observed dynamic changes in circulating and tissue hormone levels, we did not directly assess the functional consequences of altered enteroendocrine signaling on downstream metabolic outcomes such as insulin secretion, appetite regulation, or hepatic glucose handling. Additionally, although our data suggest EEC desensitization and lineage impairment with prolonged HFD, we did not employ genetic or pharmacologic tools to causally manipulate EEC differentiation or activity. Microbiota composition was not profiled, and the contribution of microbial factors to EEC remodeling remains inferential, based on prior studies [46, 59–61]. Organoid analyses were performed at the bulk level, which limits resolution of specific EEC subtypes, therefore, future studies using single-cell transcriptomics could better define shifts in enterochromaffin versus incretin-producing cells. Moreover, 5-HT co-localization with fibronectin in the liver suggests a spatial relationship with fibrotic regions, but higher-resolution imaging and quantification would be needed to confirm cellular sources and functional roles. Finally, the study was limited to male mice, and whether these findings extend to females or human MASLD/MASH remains to be determined.

## Abbreviations

2-PY: N-methyl-2-pyridone-5-carboxamide
5-HT: Serotonin
CHGA: chromogranin A
EEC: Enteroendocrine cells
HFD: High-Fat Diet
IL-6: Interleukin-6
GIP: Glucose-dependent Insulinotropic Polypeptide
GLP-1: Glucagon-like Peptide-1
GPCR: G Protein-Coupled Receptor
LCN-2: Lipocalin-2
LGR5: Leucine-rich repeat-containing G-protein coupled receptor 5
MASLD: Metabolic Dysfunction-Associated Steatotic Liver Disease
MASH: Metabolic Dysfunction-Associated Steatohepatitis
MCP-1: Monocyte Chemoattractant Protein 1
NAD: Nicotinamide Adenine Dinucleotide
PP: Pancreatic Peptide
PYY: Peptide tyrosine tyrosine
RaMP-DB: Relational database of Metabolomic Pathways
TGR5: Takeda G protein-coupled Receptor 5
TNF: Tumor Necrosis Factor

## Author’s contribution

Conception and design: JAR, ACM, MRM, EFC

Development of methodology: JAR, ACM, JGI, MRM, EFC

Acquisition of data: JAR, ACM, SSG, PP, ASR, CME, BBM, PRA, MAG, FTL, RRG, KGM, JP, SL

Analysis and interpretation of data: JAR, ACM, RRG, MRM, EFC

Writing, review, and/or Revision: JAR, ACM, SSG, PP, ASR, CME, PRA, MG, RR, KMG, JP, SL, JGI, MRM, EFC

Administrative, technical, or material support (i.e., reporting or organizing data, constructing database): JAR, ACM, MRM, EFC

Study supervision and funding acquisition: MRM and EFC

## Data availability statement

All primary data associated with this study are present in the paper or the Supplementary Materials.

## Acknowledgments

We would also like to acknowledge the Huck Institutes’ Metabolomics Core Facility (RRID:SCR_023864) for use of the OE 240 LCMS and Sergei Koshkin for helpful discussions on sample preparation and analysis.

