## Supplemental data for "Metabolomic, Lipidomic, and Enterohormone Changes in the Progression from MASLD to MASH"

Jaclyn A. Rivas *et al.*

**Contents: Materials and methods, supplementary figures legends, supplementary figures, and references.**

**Materials and methods**

**Animals.** The diet induced NASH B6 (C57BL/6NTac, Taconic Biosciences, Inc) male mice were conditioned on a modified amylin liver NASH (AMLN) diet for 16-weeks to establish MASLD and 29-weeks to establish MASH. The diet D09100310i (Research Diets, Inc) contains 40 kcal% fat, 20 Kcal% fructose, and 2% cholesterol and is irradiated. C57BL/6NTac males were put on diet D09100310i at 6 weeks of age and housed at reduced density. Control C57BL/6NTac male mice were fed the NIH-31M chow diet. All experiments were approved by the Institutional Animal Care and Use Committee of the University of New Mexico Health Sciences Center, in accordance with the National Institutes of Health guidelines for use of live animals. The University of New Mexico Health Sciences Center is accredited by the American Association of Accreditation of Laboratory Animal Care.

**Cells and tissue preparation.** Upon necropsy, all tissues were weighed, measured, sectioning out for different procedures, and flesh frozen. Blood was collected via cardiac puncture and collected in the BD P800 blood collection tube containing Proprietary Cocktail of Protease, Esterase and DPP-IV Inhibitors (BD Biosciences, catalog #: 366420) for plasma isolation. Plasma was used for serotonin (5-HT) detection via ELISA (Bio-Techne, catalog #: NBP2-68127) or hormone (Amylin; C-Peptide; Ghrelin; GIP; GLP-1 (total); Glucagon; Insulin; Leptin; PP; PYY; Resistin; and Secretin) and cytokine (IL-6, MCP-1/CCL2; TNFα) detection by a MILLIPLEX mouse metabolic hormone expanded panel multiplex assay (Millipore, catalog #: MMHE-44K). Fresh stool pellets were collected directly from the intact colon to assess lipocalin-2 levels via ELISA (Bio-Techne, catalog #: MLCN20). A section of the proximal colon and liver were stored in 10% formalin for 72 hours, then washed and stored in 70% ethanol prior for processing for tissue sectioning. Histopathological evaluation was performed on the mouse colon and liver sections using Hematoxylin and Eosin (H and E) stained slides after formalin fixation. Standard five-micron thick H and E sections obtained from paraffin embedded blocks fixed in 10% formalin were used for assessment. The histopathological evaluation was performed on the mouse liver and large intestinal sections using Hematoxylin and Eosin (H and E) stained slides. Formalin-fixed, paraffin-embedded (FFPE) sections of proximal colon were prepared at the University of New Mexico Comprehensive Cancer Center (UNMCCC) Human Tissue Repository (HTR) Core Facility. Sections (5 μm) were cut from blocks fixed in 10% neutral-buffered formalin and stained with hematoxylin and eosin (H&E) using standard protocols. The tissue slides were reviewed on an Olympus BX40 microscope at 4X, 10X, 20X and 40 X magnifications sequentially. Liver sections were evaluated using the NASH Clinical Research Network Scoring System following the guidelines provided by Liang et al [1]. Assessed histopathological parameters included - macrovesicular steatosis (+/−), microvesicular steatosis (+/−), steatosis grade (0–3), lobular inflammation (0–3), hepatocellular hypertrophy (0–2), and fibrosis stage (0–4). Additional fibrosis staging was performed using the Ishak scoring system. The degree of intestinal inflammation scoring was performed based on the guidelines provided by Erben et al [2]. Colon sections were scored for histopathological features of colitis based on established criteria, including: submucosal edema (0–3), polymorphonuclear leukocyte (PMN) infiltration (0–3), goblet cell depletion (0–3), and epithelial integrity (0–3).

**Generation of intestinal organoids**. Intestinal organoid media was comprised of Advanced Dulbecco's modified Eagle medium/Ham's F-12 (ThermoFisher, Waltham, MA), 100 U/mL penicillin/streptomycin (Quality Biological, Gaithersburg, MD), WNT surrogate-Fc fusion protein (ImmunoPrecise Antibodies Ltd, Utrecht, The Netherlands) 15% v/v R-spondin1 conditioned medium (cell line kindly provided by Calvin Kuo, Stanford University), 10% v/v Noggin conditioned medium (cell line kindly provided by Gijs van den Brink, Tytgat Institute for Liver and Intestinal Research), 1X B27 supplement (ThermoFisher), 10 mM HEPES (ThermoFisher), 1X GlutaMAX (ThermoFisher), 1mM N-acetylcysteine (MilliporeSigma), 50 ng/mL human epidermal growth factor (ThermoFisher), 10 nM [Leu-15] gastrin (AnaSpec, Fremont, CA), 500 nM A83-01 (Tocris, Bristol, United Kingdom), 10 μM SB202190 (MilliporeSigma), 100 mg/mL primocin (InvivoGen, San Diego, CA). Base media, for differentiation of organoids, had the same composition of media but lacked WNT surrogate Fc fusion protein, Rspo-1 and SB202190. Colonic crypt isolation and colonic organoid generation were prepared as previously reported [3-6]. Isolated crypts were resuspended in Matrigel (Corning) and 25 μL droplets were plated in a 24-well plate (Corning). After polymerization at 37°C, 500μL of organoid expansion media was added for 2 d. After 2 d, the organoid expansion media was replaced every other day. Colonic organoids were passaged every 3-7 d by harvesting in Cultrex Organoid Harvesting Solution (Bio-Techne, Minneapolis, MN) at 4°C with shaking for 45 min, as previously described [5, 6]. All colonic organoids cultures were maintained at 37°C and 5% CO2. Unless noted, colonoid lines have been passaged >30 times. Colonic organoids were harvested from Matrigel using Cultrex Organoid Harvesting Solution as previously described.

**Immunofluorescence, Microscopy, and image analysis.** Formalin-fixed, paraffin-embedded mouse (intestinal or liver) sections were baked at 60 °C for 90 min and cooled to room temperature. Slides were deparaffinized with three washes in xylene (15 min each), followed by rehydration through a graded ethanol series (three washes in 100% ethanol for 5 min each, two washes in 95% ethanol for 5 min each, one wash in 70% ethanol for 5 min, and one wash in 50% ethanol for 5 min) before rinsing in running distilled water for 5–10 min. Sections were further rehydrated by two washes in 1X PBS containing 0.1% Tween-20 (10 min each). After encircling the tissue with a hydrophobic barrier pen, samples were incubated in blocking buffer (0.1% saponin, 2% BSA, and 15% FBS in 1X PBS). Primary antibodies were applied overnight at 4 °C in a humidified chamber, with 100 µl of antibody solution prepared in 1X PBS. CHGA (Abcam) and 5-HT (Abcam) were prepared at 1:200 dilutions, while fibronectin (Invitrogen) was prepared at 1:500. After three washes in 1X PBS, 100 µl of secondary antibody at a 1:200 dilution in 1X PBS plus DAPI (ThermoFisher; 1 mg/ml) was added for 1 h at room temperature**.** After three final washes in 1X PBS**,** 2.5 M fructose–60% glycerol stock was added, and a number 1 coverslip was placed on top of the tissue. Confocal imaging of fluorescently labeled tissue sections was performed at the UNM Comprehensive Cancer Center using the Olympus iX83 – Yokogawa Spinning Disk Confocal Microscope and images were processed using the Olympus OlyVIA 4.2 software. Imaging of H&E-stained tissue sections was performed using the Evident APEXVIEW APX100 Benchtop Fluorescence Microscope. Acquisition parameters in the cellSens APEX software were standardized across all samples, and images were exported as high-resolution TIFF files. All primary and secondary antibodies used for confocal staining are listed in Table S5.

**RNA Isolation, Quantification, and RT-qPCR.** RNA isolation was performed using the RNA Purelink Minikit (Invitrogen) according to the manufacturer's protocols on colonic organoids which were harvested from Matrigel using Cultrex Organoid Harvesting Solution as previously described [5]. . RNA was quantified on Nanodrop2000 and all samples yielded a 260/280 of 2 ± 0.15. cDNA synthesis was performed using SuperScript IV VILO Master Mix with eZDnase enzyme(ThermoFisher). Reverse transcription reaction utilized the viiA7 Thermal Cycler (ThermoFisher and RT-qPCR runs utilized Taqman MasterMix (ThermoFisher) using the QuantStudio 7. RT-qPCR primers and reagents used are listed in Table S6.

**Methods for Metabolomics.** Metabolite Extraction: Tissues samples (Liver, Colon, Stool) were weighed, homogenized with liquid nitrogen in a cryomill (Retsch) at 25Hz for 45 seconds, before extracting with 40:40:20 acetonitrile: methanol: water, with a volume of 40μl solvent per 1mg of sample [7]. The samples were vortexed for 15 seconds, incubated on ice for 10 minutes, and centrifuged at 16,000 g for 30 minutes. The supernatants transferred to new Eppendorf tubes and centrifuged at 16,000 g for 30 minutes. The top 50μl of the supernatant was used for LC-MS analysis. LC-MS: Extracts were analyzed within 24 hours by liquid chromatography coupled to a mass spectrometer (LC-MS). The LC–MS method was based on hydrophilic interaction chromatography (HILIC) coupled to the Orbitrap Exploris 240 mass spectrometer (Thermo Scientific) [8]. The LC separation was performed on a XBridge BEH Amide column (2.1 x 150 mm, 3.5 $\mu$m particle size, Waters, Milford, MA). Solvent A is 95%: 5% H2O: acetonitrile with 20 mM ammonium acetate and 20mM ammonium hydroxide, and solvent B is 90%: 10% acetonitrile: H2O with 20 mM ammonium acetate and 20mM ammonium hydroxide. The gradient was 0 min, 90% B; 2 min, 90% B; 3 min, 75% B; 5 min, 75% B; 6 min, 75% B; 7 min, 75% B; 8 min, 70% B; 9 min, 70% B; 10 min, 50% B; 12 min, 50% B; 13 min, 25% B; 14min, 25% B; 16 min, 0% B; 18 min, 0% B; 20 min, 0% B; 21 min, 90% B; 25 min, 90% B. The following parameters were maintained during the LC analysis: flow rate 150 mL/min, column temperature 25 °C, injection volume 5 μL and autosampler temperature was 5 °C. For the detection of metabolites, the mass spectrometer was operated in both negative and positive ion mode. The following parameters were maintained during the MS analysis: resolution of 180,000 at m/z 200, automatic gain control (AGC) target at 3e6, maximum injection time of 30 ms and scan range of m/z 70-1000. Raw LC/MS data were converted to mzXML format using the command line “msconvert” utility [9]. Data was analyzed via the EL-MAVEN software version 12.

**Methods for Lipidomics.** Tissue homogenization and extraction for lipids: Tissue samples were homogenized using a Retsch CryoMill. The homogenates were mixed with 1 mL of Extraction Buffer containing IPA/H2O/Ethyl Acetate (30:10:60, v/v/v) and Avanti Lipidomix Internal Standard (diluted 1:1000) (Avanti Polar Lipids, Inc. Alabaster, AL). Samples were vortexed and transferred to bead mill tubes for homogenization using a VWR Bead Mill at 6000 g for 30 seconds, repeated twice. The samples were sonicated for 5 minutes and then centrifuged for 5 minutes at 15,000 g/4°C. The upper phase was transferred to a new tube and kept at 4°C. Another 1 mL of Extraction Buffer (30:10:60, v/v/v) was added to the tissue pellet-containing tube to re-extract the tissues. The samples were vortexed, homogenized, sonicated, and centrifuged. The supernatants from both extractions were combined, and the organic phase was dried under liquid nitrogen gas. Sample reconstitution for lipids: The dried tissue samples were reconstituted in 300 μL of

Solvent A (IPA/ACN/H2O, 45:35:20, v/v/v). After brief vortexing, the samples were sonicated for 7 minutes and centrifuged at 15,000 g for 10 minutes at 4°C. The supernatants were transferred to clean tubes and centrifuged again for 5 minutes at 15,000 g at 4°C to remove any remaining particulates. For LC-MS lipidomic analysis, 60 µL of the sample extracts were transferred to mass spectrometry vials. LC-MS analysis for lipids: Sample analysis was performed within 24 hours after extraction using a Vanquish UHPLC system coupled with an Orbitrap Exploris 240™ mass spectrometer equipped with a H-ESI™ ion source (all Thermo Fisher Scientific). A Waters (Milford, MA) CSH C18 column (1.0 × 150 mm × 1.7 µm particle size) was used. Solvent A consisted of ACN:H2O (60:40; v/v) with 10 mM Ammonium formate and 0.1% formic acid, while solvent B contained IPA:ACN (95:5; v/v) with 10 mM Ammonium formate and 0.1% formic acid. The mobile phase flow rate was set at 0.11 mL/min, and the column temperature was maintained at 65 °C. The gradient for solvent B was as follows: 0 min 15% (B), 0–2 min 30% (B), 2–2.5 min 48% (B), 2.5–11 min 82% (B), 11–11.01 min 99% (B), 11.01–12.95 min 99% (B), 12.95–13 min 15% (B), and 13–15 min 15% (B). Ion source spray voltages were set at 4,000 V and 3,000 V in positive and negative mode, respectively. Full scan mass spectrometry was conducted with a scan range from 200 to 1000 m/z, and AcquireX mode was utilized with a stepped collision energy of 30% with a 5% spread for fragment ion MS/MS scan. Statistical analysis for lipidomics: After normalization, all data were analyzed in R using the *lipidr* package [10]. Data sets for each tissue type were analyzed independently. Data was log transformed and further processed before analysis. For lipids with multiple readings across replicates, only the sets of readings with the highest values were used. In a few instances (2–3), for each tissue type, all samples had identical measurements. As this was likely due to technical errors, these samples were not used. Lastly, to be compatible with *lipidr*, lipid names had to be modified to fit a standard "CLS xx:x/yy:y" naming scheme where CLS refers to the abbreviated lipid class and xx:x and yy:y refer to the first and second chains. The lipid composition of samples was compared after processing using the "de_analysis" function from *lipidr* with default settings. Here, *lipidr* uses moderated *t*-tests to identify significant differences in lipids between sample types within a tissue type. Significantly different lipids were those with adjusted $p$-values < 0.05 (*note*: $p$-values were adjusted to correct for multiple comparisons using a false discovery rate procedure) and log fold changes greater than 1 or less than −1. These results were used to perform a lipid set enrichment analysis using the "lsea" function in which entries were ranked by fold change; only classes with at least four associated lipids were considered, and 100,000 permutations were run. Here, the method *lipdr* used is based on the commonly used gene set enrichment analysis approach previously outlined [11]. Briefly, lipid class and chain length categories were determined from annotations extracted from lipid names in the data set, and lipids were ranked by fold change. A permutation algorithm was used to calculate enrichment scores and *p*-values for each lipid set. Sets with adjusted *p*-values < 0.05 were defined as significantly enriched. Lastly, heatmaps were generated for significantly enriched lipid classes using the "plot_heatmap".

**Statistical Analysis.** Metabolomic data from colon, liver, and stool samples were analyzed relative to untreated control mice. Sample groups included: colon (16-week control n = 7; 16-week HFD n = 8; 29-week control n = 8; 29-week HFD n = 8), liver (16-week control n = 8; 16-week HFD n = 8; 29-week control n = 8; 29-week HFD n = 8), and stool (16-week control n = 7; 16-week HFD n = 5; 29-week control n = 7; 29-week HFD n = 5). All analyses were performed using MetaboAnalyst 6.0 (www.metaboanalyst.ca). Data were normalized by sum, log-transformed, and mean-centered prior to analysis. A false discovery rate (FDR) correction was applied where appropriate, with a significance threshold of $p = 0.05$. A 2.0-fold change cutoff was applied under the assumption of equal variance. Volcano plots were generated to visualize significantly altered metabolites, defined by both fold-change (≥2.0) and statistical significance ($p < 0.05$). Quantitative enrichment analysis was performed using KEGG IDs for metabolite annotation. Data were normalized by sum, log2-transformed, and auto-scaled. Enrichment was conducted against metabolite set libraries derived from RaMP-DB (pathway-based) and disease signatures (blood or feces). Only metabolite sets containing at least five entries were included, and all available compounds within each selected library were used in the analysis. To evaluate the association between cholic acid and enteroendocrine hormone secretion, linear regression analyses were performed in GraphPad Prism 10 (San Diego, CA). Data from the 16-week colon groups (control $n = 7$; HFD $n = 8$) were analyzed separately for GIP and GLP-1. A simple linear regression model was fitted to assess the relationship, with regression coefficients, 95% confidence intervals, coefficient of determination ($R^2$), and associated *p*-values reported. Significance was defined as $p < 0.05$. Scatter plots with best-fit regression lines and 95% confidence bands were generated to visualize the associations.

**Supplementary Figures and Figure Legends**

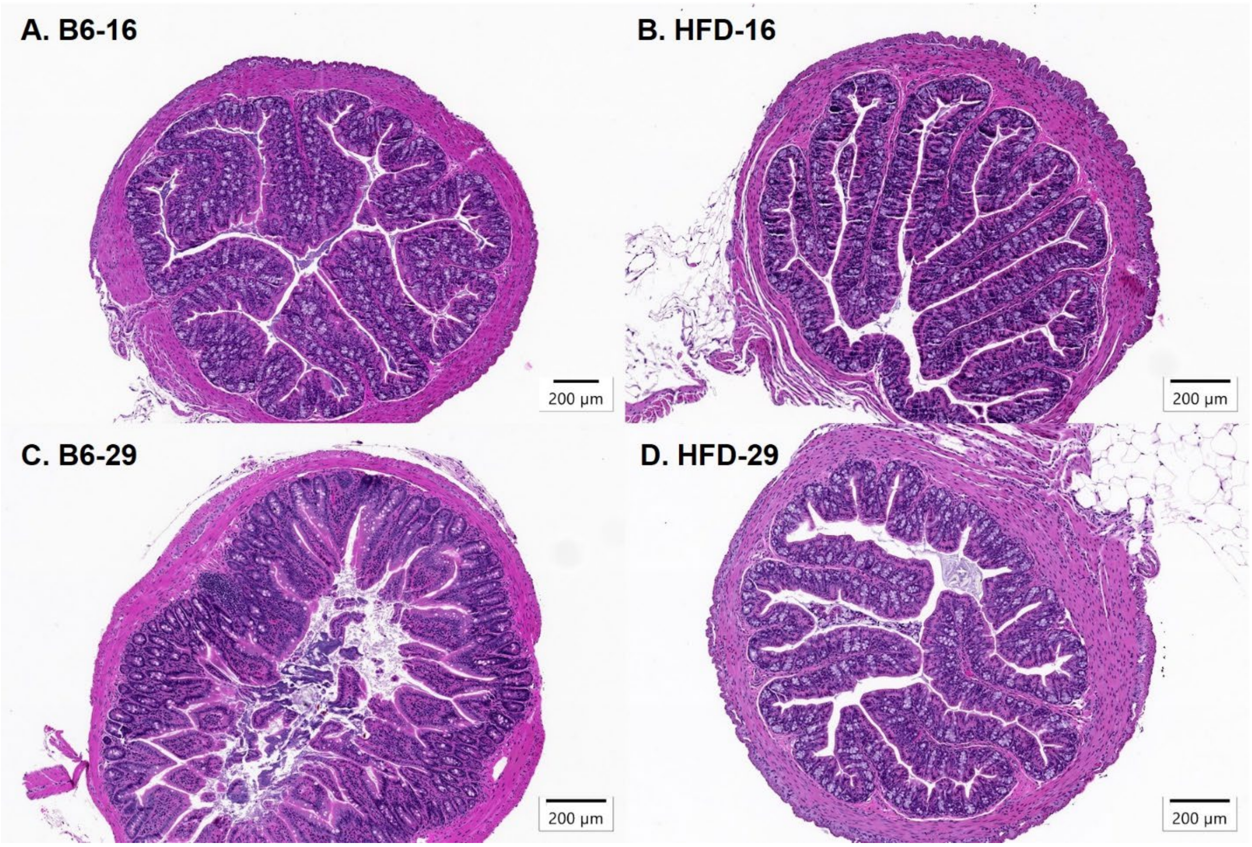


**Supplementary Figure 1. Histological assessment of colons.** Representative H&E-stained colon sections from (**A**) B6-16, (**B**) HFD-16, (**C**) B6-29 and (**D**) HFD-29 mice; histopathologic scoring summarized in Table S2. n = 8 mice per group.

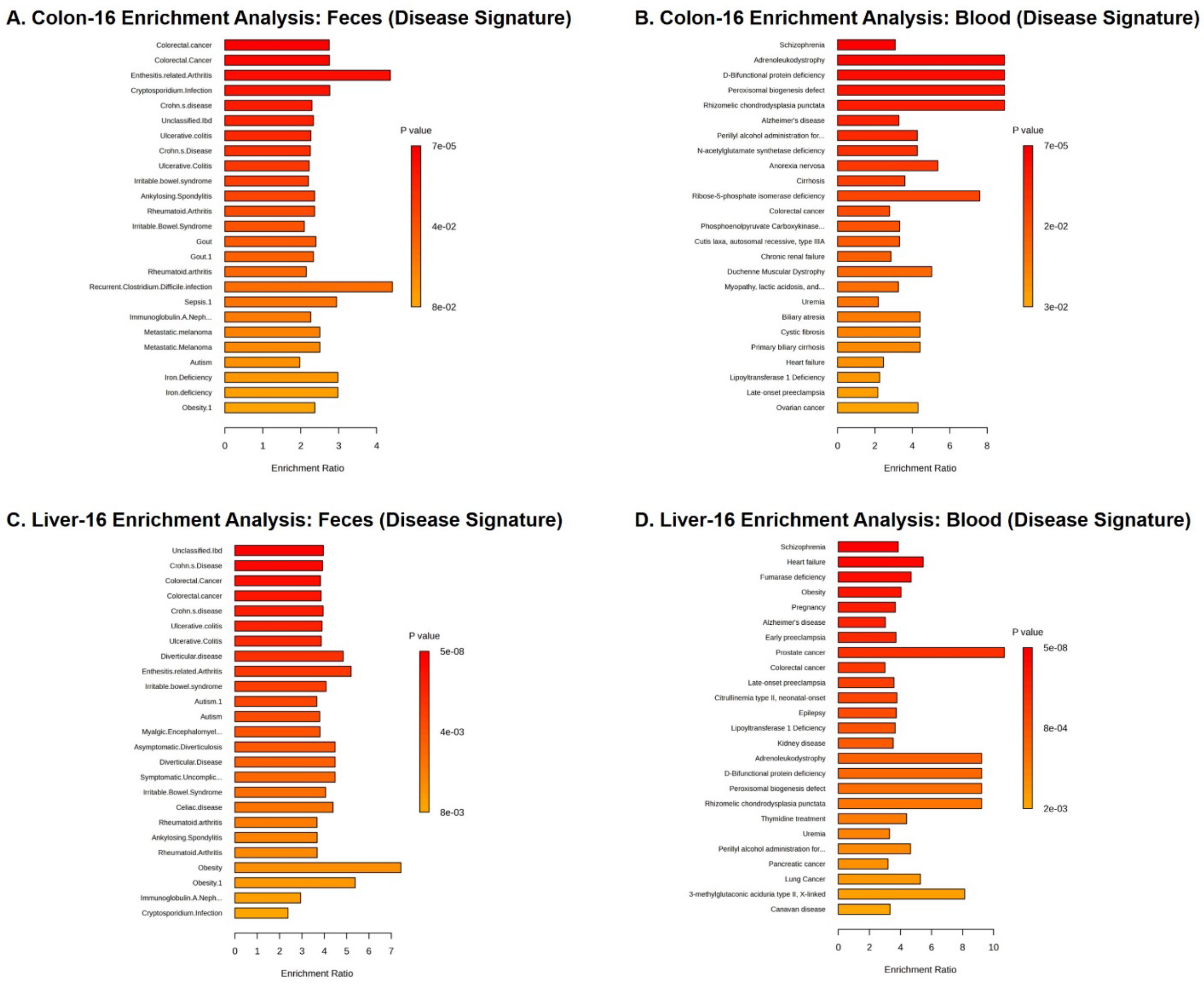


**Supplementary Figure 2. Disease signature enrichment of colon and liver metabolites from HFD-16 mice.** To evaluate disease relevance, differentially abundant colonic and hepatic metabolites from HFD-fed mice (16 weeks) in comparison to controls were mapped to known human fecal and blood disease-associated metabolite signatures using RaMP-DB. (**A**, **B**) Enrichment analysis of colon-derived metabolites revealed significant overlap with fecal (**A**) and blood (**B**) disease signatures. (**C**, **D**) Similarly, liver-derived metabolite signatures were significantly associated with fecal (**C**) and blood (**D**) disease profiles. Color indicates p-value, and bar length represents enrichment ratio.

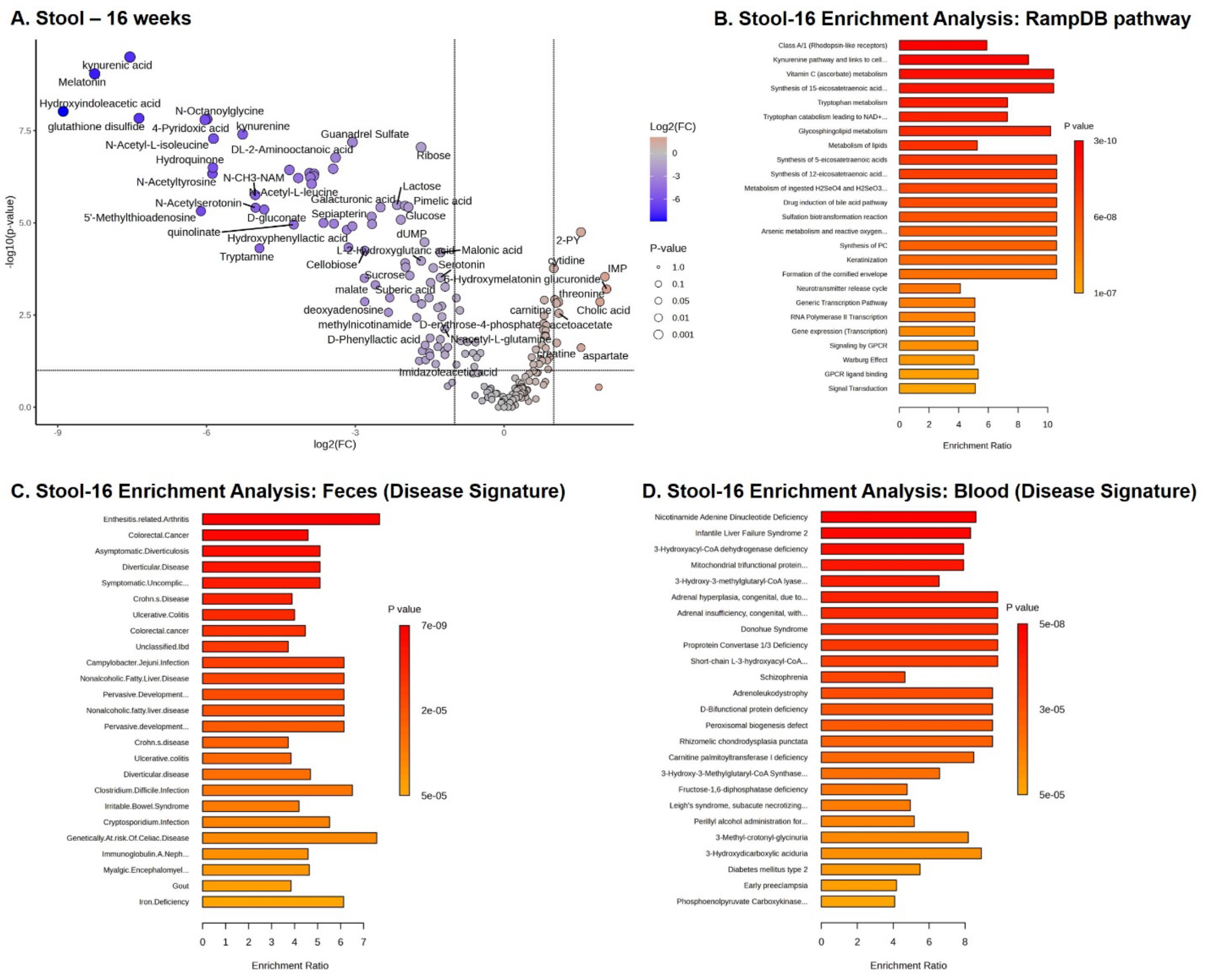


**Supplementary Figure 3. Metalobolomic alterations and Disease signature enrichment of stool metabolites from 16-week groups.** (**A**) Volcano plots show significantly altered metabolites in the stool of mice fed a high-fat diet (HFD-16) compared to age-matched controls on a normal chow diet. $Log_2$ fold change (x-axis) is plotted against $–log_{10}$ (p-value) (y-axis), with dot size indicating significance level and color indicating directionality of change. (**B**) Metabolite enrichment analysis using RaMP-DB highlights pathway-level changes in stool. (**C**, **D**) Enrichment analysis of colon-derived metabolites revealed significant overlap with fecal (**C**) and blood (**D**) disease signatures. Color indicates p-value, and bar length represents enrichment ratio.

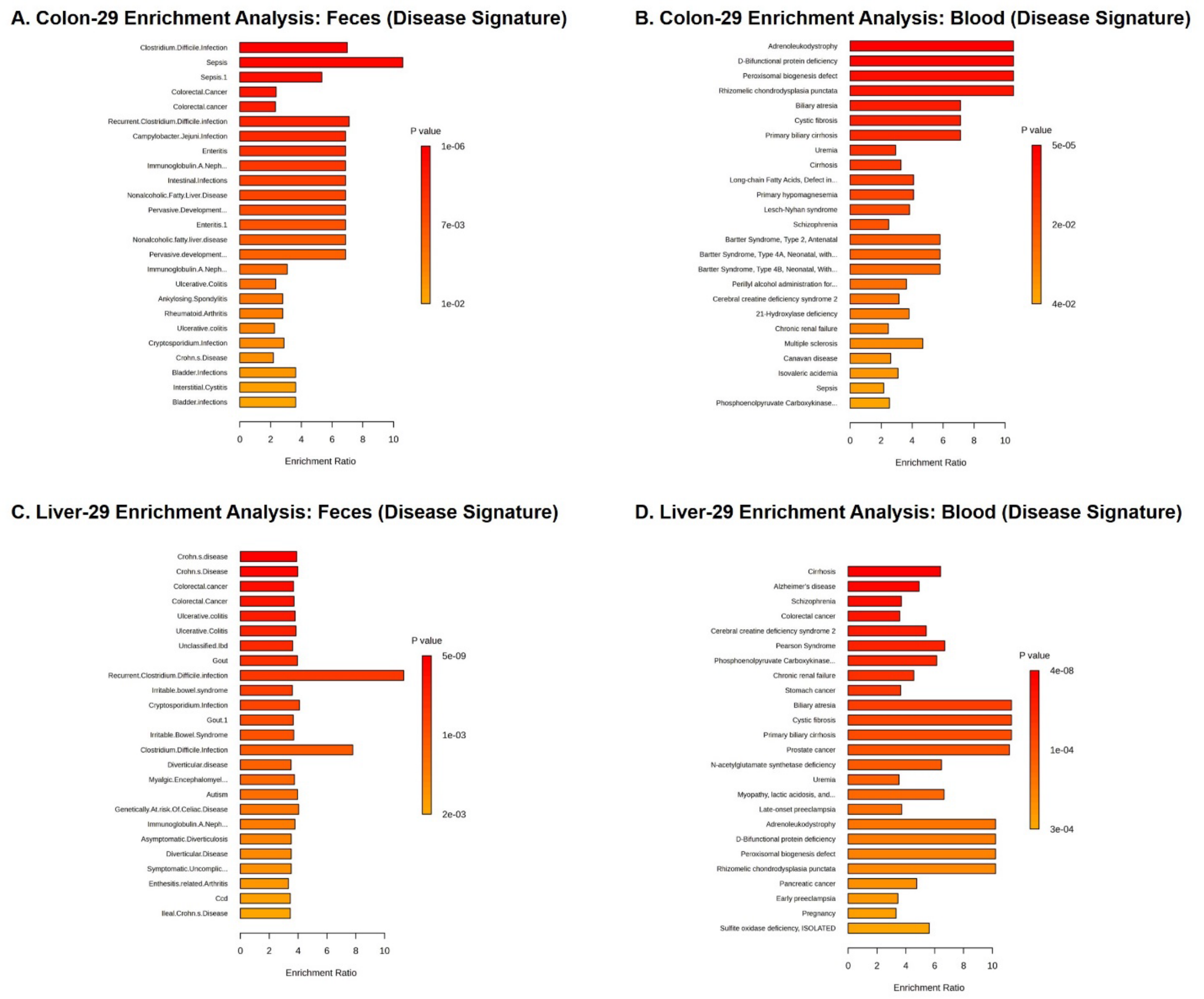


**Supplementary Figure 4. Disease signature enrichment of colon and liver metabolites from HFD-29 mice.** To evaluate disease relevance, differentially abundant colonic and hepatic metabolites from HFD-fed mice (29 weeks) in comparison to controls were mapped to known human fecal and blood disease-associated metabolite signatures using RaMP-DB. (**A**, **B**) Enrichment analysis of colon-derived metabolites revealed significant overlap with fecal (**A**) and blood (**B**) disease signatures. (**C**, **D**) Similarly, liver-derived metabolite signatures were significantly associated with fecal (**C**) and blood (**D**) disease profiles. Color indicates p-value, and bar length represents enrichment ratio.

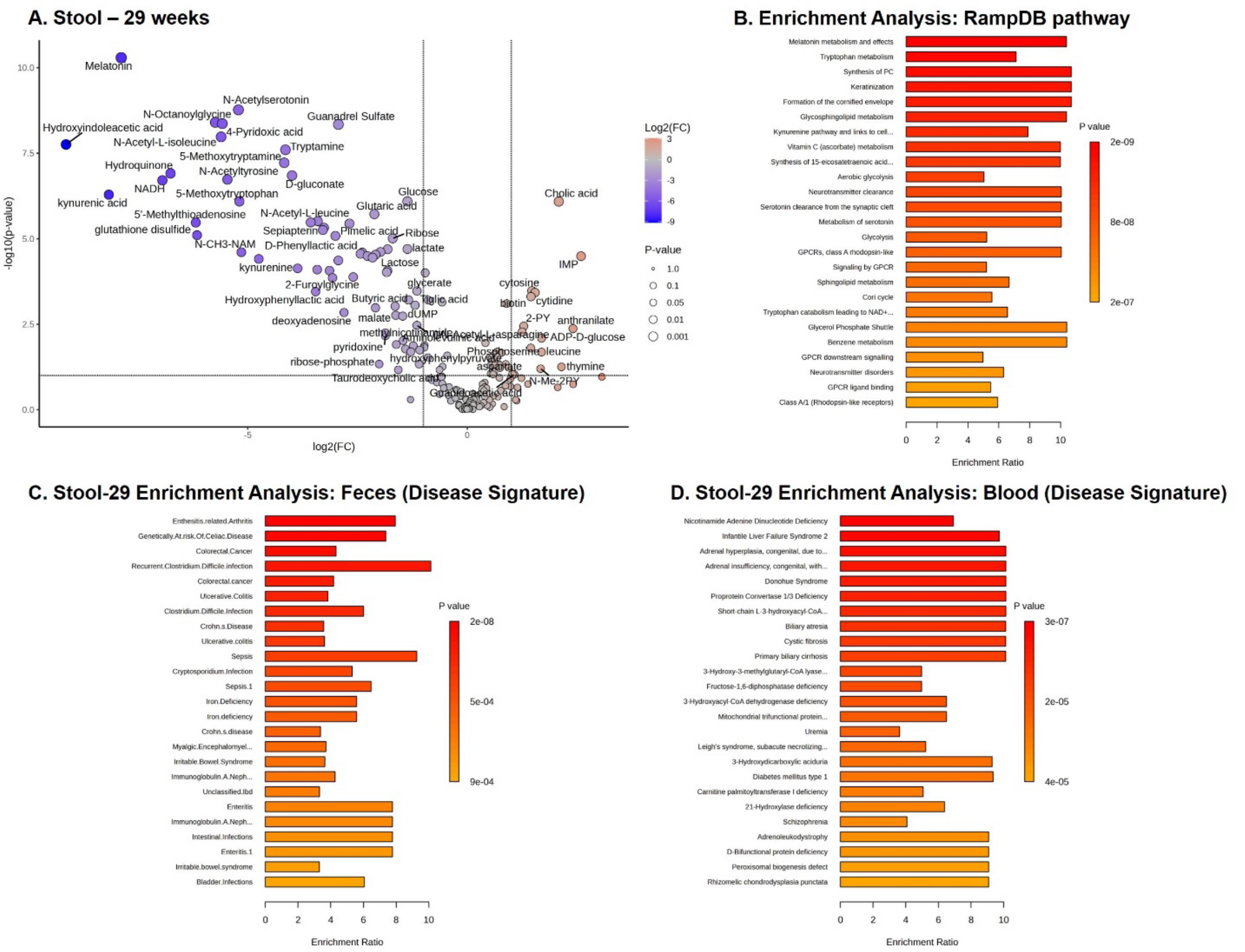


**Supplementary Figure 5. Metalobolomic alterations and Disease signature enrichment of stool metabolites from 29-week groups.** (**A**) Volcano plots show significantly altered metabolites in the stool of mice fed a high-fat diet (HFD-29) compared to age-matched controls on a normal chow diet. $Log_2$ fold change (x-axis) is plotted against $-log_{10}$ (p-value) (y-axis), with dot size indicating significance level and color indicating directionality of change. (**B**) Metabolite enrichment analysis using RaMP-DB highlights pathway-level changes in stool. Enrichment analysis of colon-derived metabolites revealed significant overlap with fecal (**C**) and blood (**D**) disease signatures. Color indicates p-value, and bar length represents enrichment ratio.

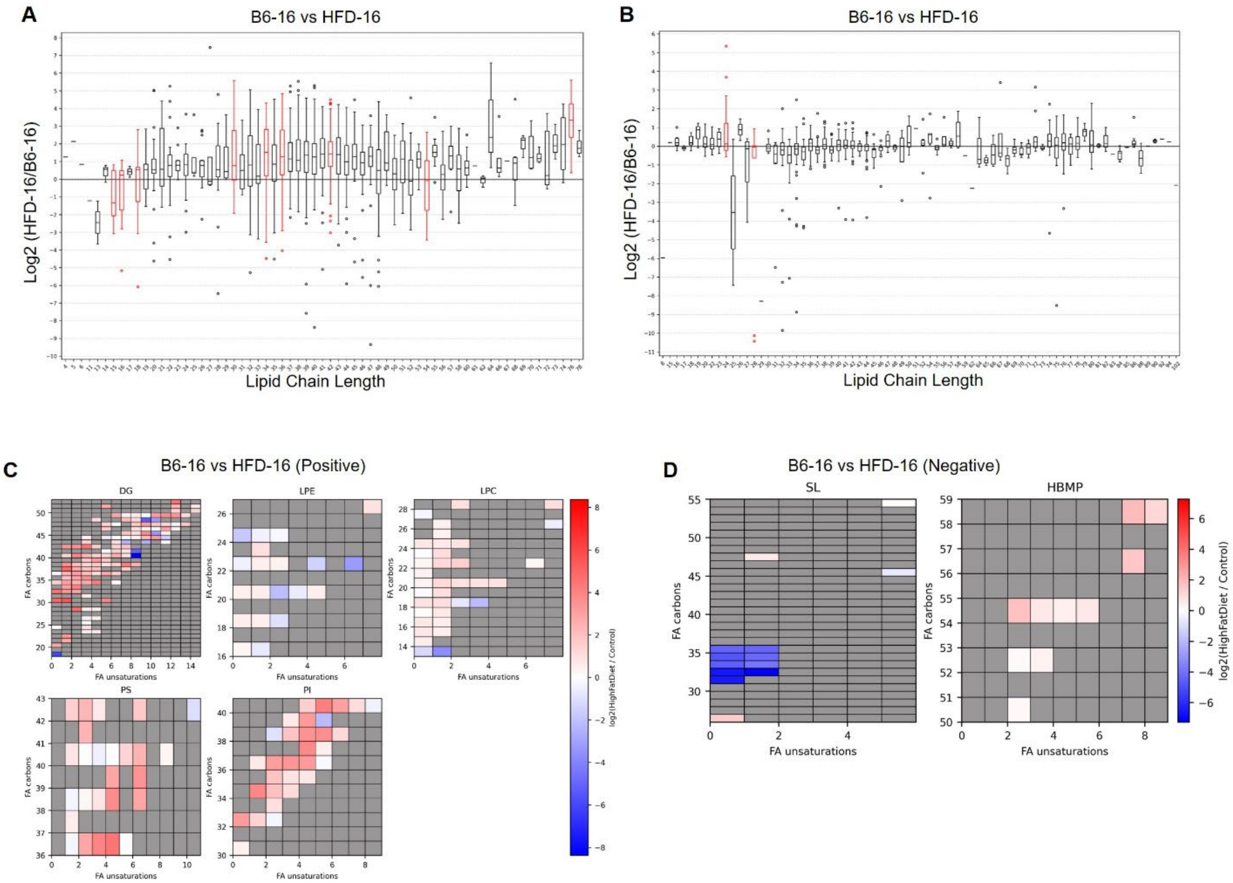


**Supplementary Figure 6. Colon lipid chain-length enrichment and species alterations in HFD-fed mice at 16-weeks**. (A) The box plot shows the log2-fold change in lipid chain-length enrichment in the colon of HFD-fed mice (HFD-16), with age-matched controls on normal chow diet as the reference (positive ionization mode). Red boxes are significant and black boxes are not significant. (B)The box plot shows the log2-fold change in lipid chain-length enrichment in the colon of mice fed a high-fat diet (HFD-16), with age-matched controls on normal chow diet as the reference (negative ionization mode). Red boxes are significant and black boxes are not significant. (C) Individual heat maps of significantly altered lipid abundance by chain length and saturation. Heat map color depicts log2FoldChange where a positive log2FoldChange indicates increased abundance in the colon of mice fed a high-fat diet (HFD-16), and a negative log2FoldChange indicates decreased abundance relative to age-matched controls on normal chow diet (positive ionization mode). (D) Individual heat maps of significantly altered lipid abundance by chain length and saturation. Heat map color depicts log2FoldChange where a positive log2FoldChange indicates increased abundance in the colon of mice fed a high-fat diet (HFD-16), and a negative log2FoldChange indicates decreased abundance relative to age-matched controls on normal chow diet (negative ionization mode). Table S4 details lipid abbreviation.

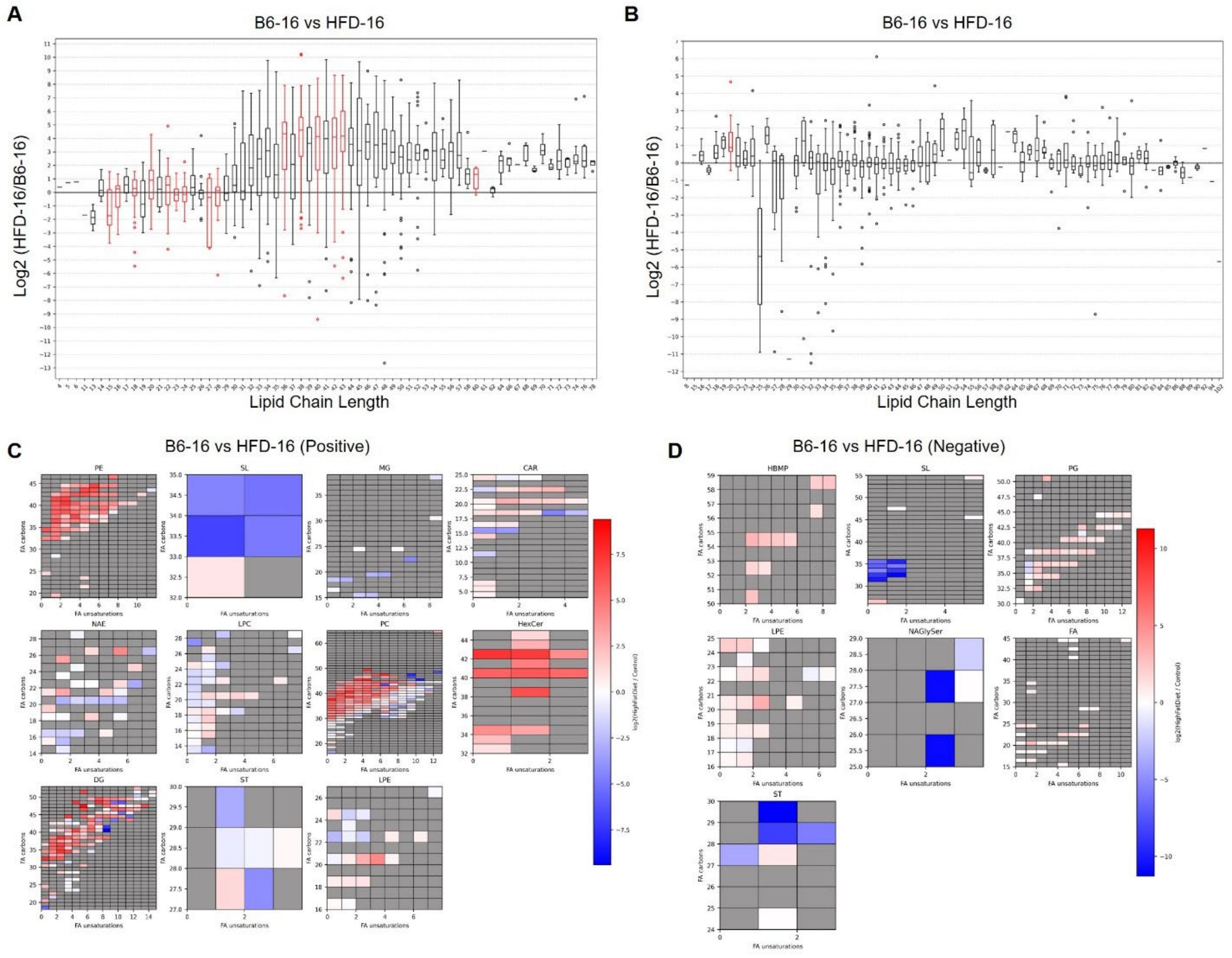


**Supplemental Figure S7. Liver lipid chain-length and saturation alterations in HFD-fed mice compared to controls at 16-weeks.** (A) The box plot shows the log2-fold change in lipid chain-length enrichment in the liver of HFD-fed mice (HFD-16), with age-matched controls on normal chow diet as the reference (positive ionization mode). Red boxes are significant and black boxes are not significant. (B) The box plot shows the log2-fold change in lipid chain-length enrichment in the liver of mice fed a high-fat diet (HFD-16), with age-matched controls on normal chow diet as the reference (negative ionization mode). Red boxes are significant and black boxes are not significant. (C) Individual heat maps of significantly altered lipid abundance by chain length and saturation. Heat map color depicts log2FoldChange where a positive log2FoldChange indicates increased abundance in the liver of mice fed a high-fat diet (HFD-16), and a negative log2FoldChange indicates decreased abundance relative to age-matched controls on normal chow diet (positive ionization mode). (D) Individual heat maps of significantly altered lipid abundance by chain length and saturation. Heat map color depicts log2FoldChange where a positive log2FoldChange indicates increased abundance in the liver of mice fed a high-fat diet

(HFD-16), and a negative log2FoldChange indicates decreased abundance relative to age-matched controls on normal chow diet (negative ionization mode). Table S4 details lipid abbreviation.

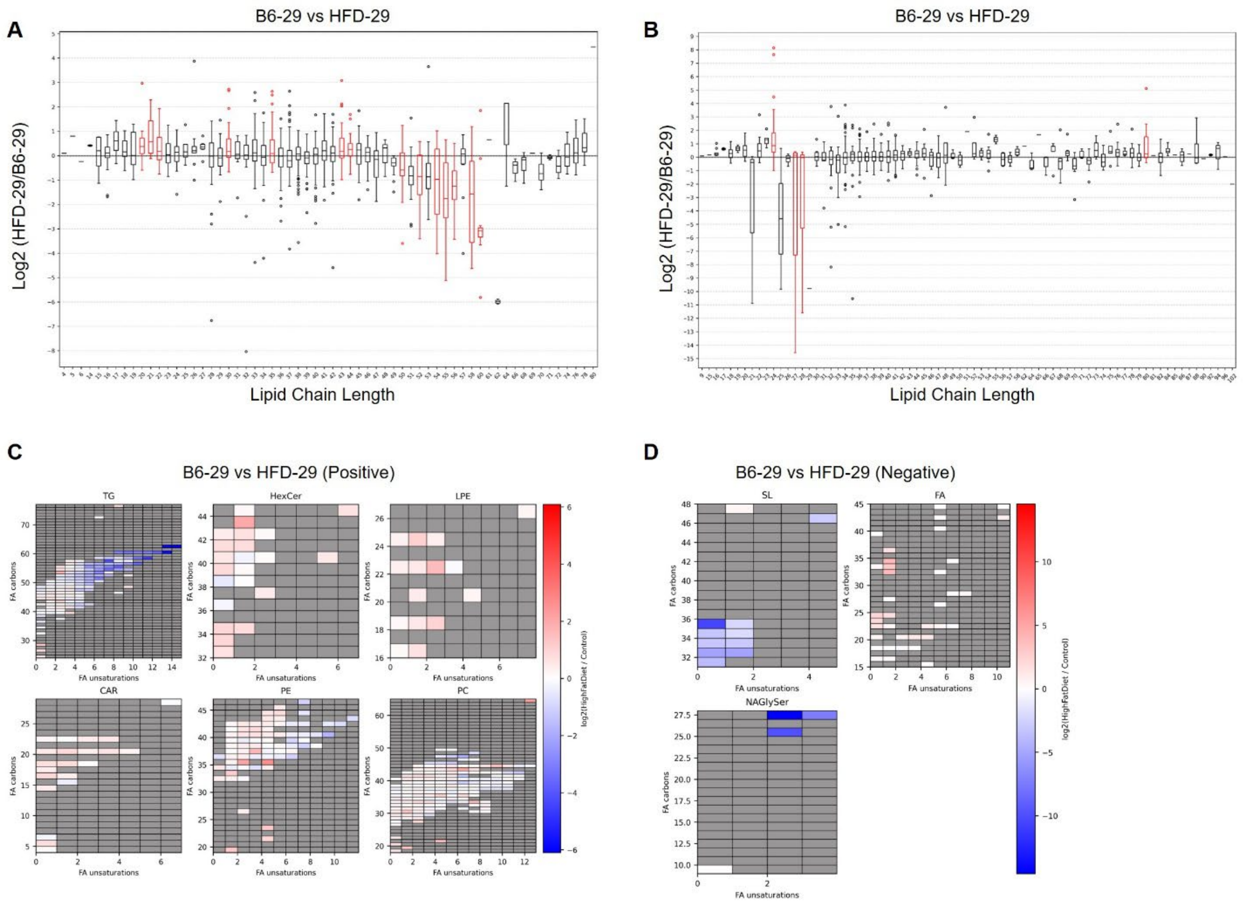


**Supplementary Figure 8. Colon lipid chain-length enrichment and species alterations in HFD-fed mice at 29-weeks**. (A) The box plot shows the log2-fold change in lipid chain-length enrichment in the colon of HFD-fed mice (HFD-29), with age-matched controls on normal chow diet as the reference (positive ionization mode). Red boxes are significant and black boxes are not significant. (B) The box plot shows the log2-fold change in lipid chain-length enrichment in the colon of mice fed a high-fat diet (HFD-29), with age-matched controls on normal chow diet as the reference (negative ionization mode). Red boxes are significant and black boxes are not significant. (C) Individual heat maps of significantly altered lipid abundance by chain length and saturation. Heat map color depicts log2FoldChange where a positive log2FoldChange indicates increased abundance in the colon of mice fed a high-fat diet (HFD-29), and a negative log2FoldChange indicates decreased abundance relative to age-matched controls on normal chow diet (positive ionization mode). (D) Individual heat maps of significantly altered lipid abundance by chain length and saturation. Heat map color depicts log2FoldChange where a positive log2FoldChange indicates increased abundance in the colon of mice fed a high-fat diet (HFD-29), and a negative log2FoldChange indicates decreased abundance relative to age-matched controls on normal chow diet (negative ionization mode). Table S4 details lipid abbreviation.

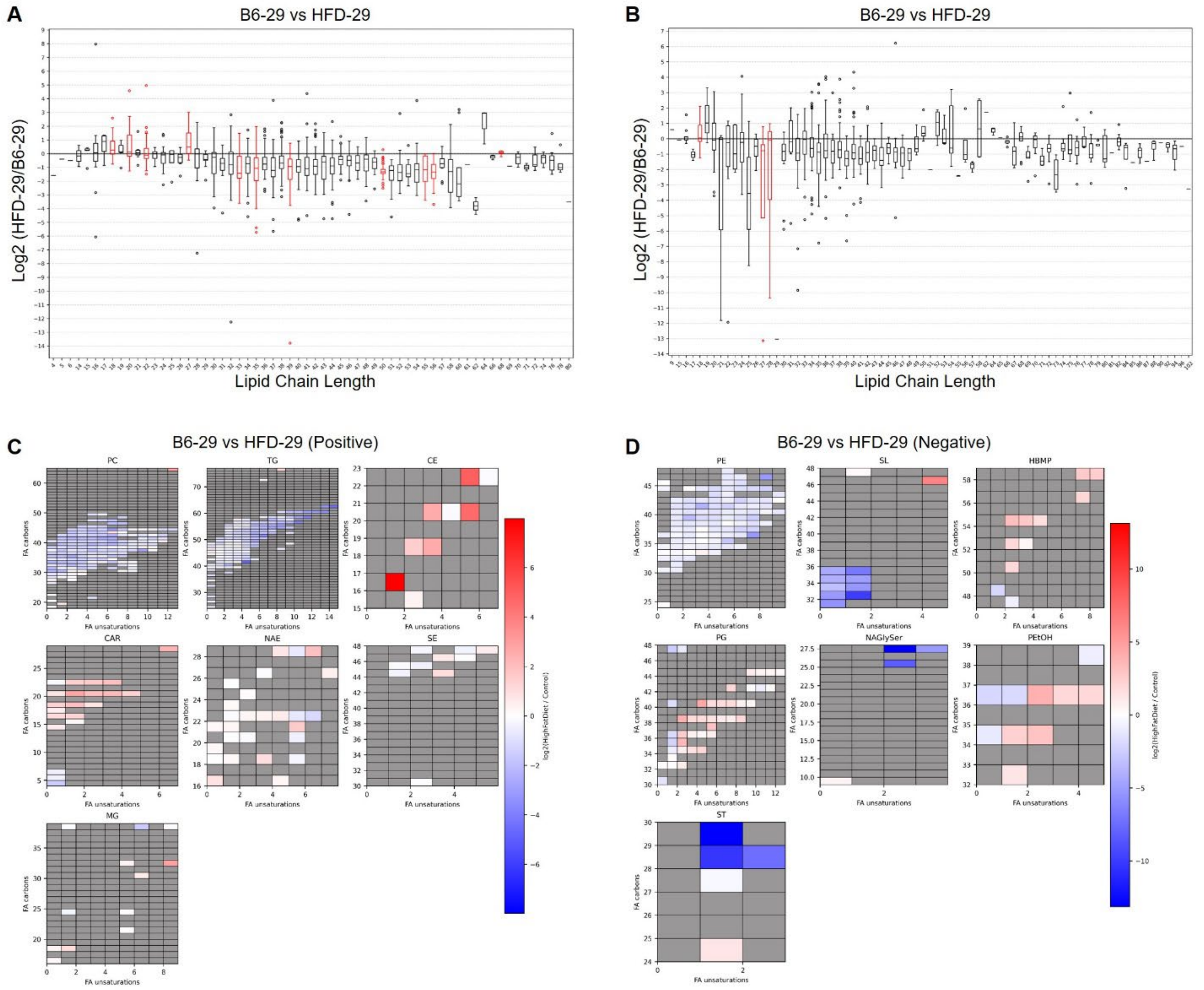


**Supplementary Figure 9. Liver lipid chain-length and saturation alterations in HFD-fed mice compared to controls at 29-weeks.** (A) The box plot shows the log2-fold change in lipid chain-length enrichment in the liver of HFD-fed mice (HFD-29), with age-matched controls on normal chow diet as the reference (positive ionization mode). Red boxes are significant and black boxes are not significant. (B) The box plot shows the log2-fold change in lipid chain-length enrichment in the liver of mice fed a high-fat diet (HFD-29), with age-matched controls on normal chow diet as the reference (negative ionization mode). Red boxes are significant and black boxes are not significant. (C) Individual heat maps of significantly altered lipid abundance by chain length and saturation. Heat map color depicts log2FoldChange where a positive log2FoldChange indicates increased abundance in the liver of mice fed a high-fat diet (HFD-29), and a negative log2FoldChange indicates decreased abundance relative to age-matched controls on normal chow diet (positive ionization mode). (D) Individual heat maps of significantly altered lipid abundance by chain length and saturation. Heat map color depicts log2FoldChange where a positive log2FoldChange indicates increased abundance in the liver of mice fed a high-fat diet (HFD-29), and a negative log2FoldChange indicates decreased abundance relative to age-matched controls on normal chow diet (negative ionization mode). Table S4 details lipid abbreviation.

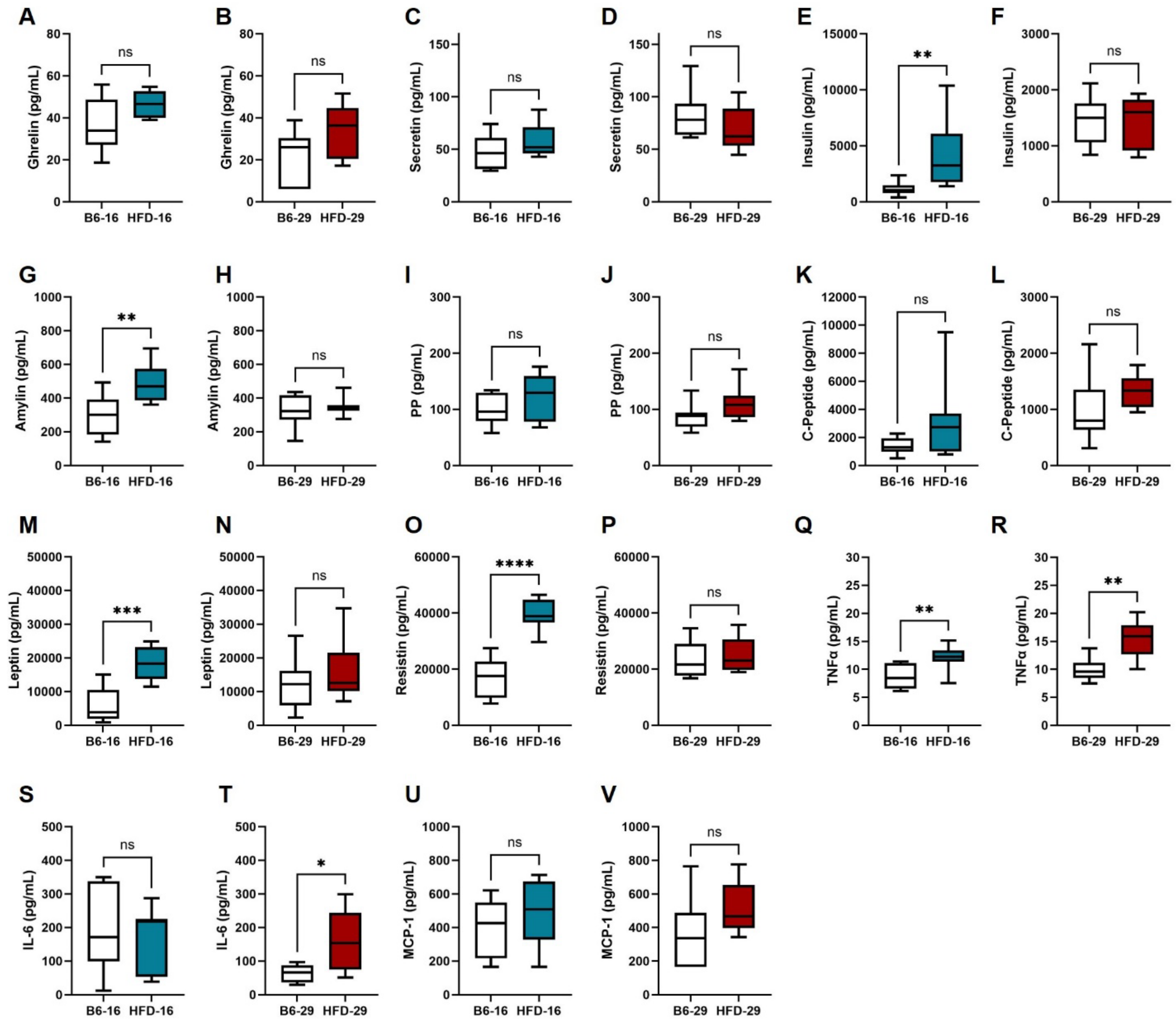


**Supplementary Figure 10. Serum cytokine and hormone levels in MASLD and MASH mice.** (**A-V**) Circulating levels of cytokines and hormones were measured in serum from mice fed a HFD for 16 weeks (HFD-16) or 29 weeks (HFD-29) and compared to age-matched controls on a normal diet (B6-16 or B6-29). (**A, B**) Ghrelin; (**C, D**) Secretin; (**E, F**) Insulin; (**G, H**) Amylin; (**I, J**) PP; (**K, L**) C-Peptide; (**M, N**) Leptin; (**O, P**) Resistin; (**Q, R**) TNFα; (**S, T**) IL-6; (**U, V**) MCP-1. n = 8 mice per group. Data analyzed using Student's t-test. *P < 0.05, **P < 0.005, ***P < 0.0005, ***P < 0.0001, NS = not significant.

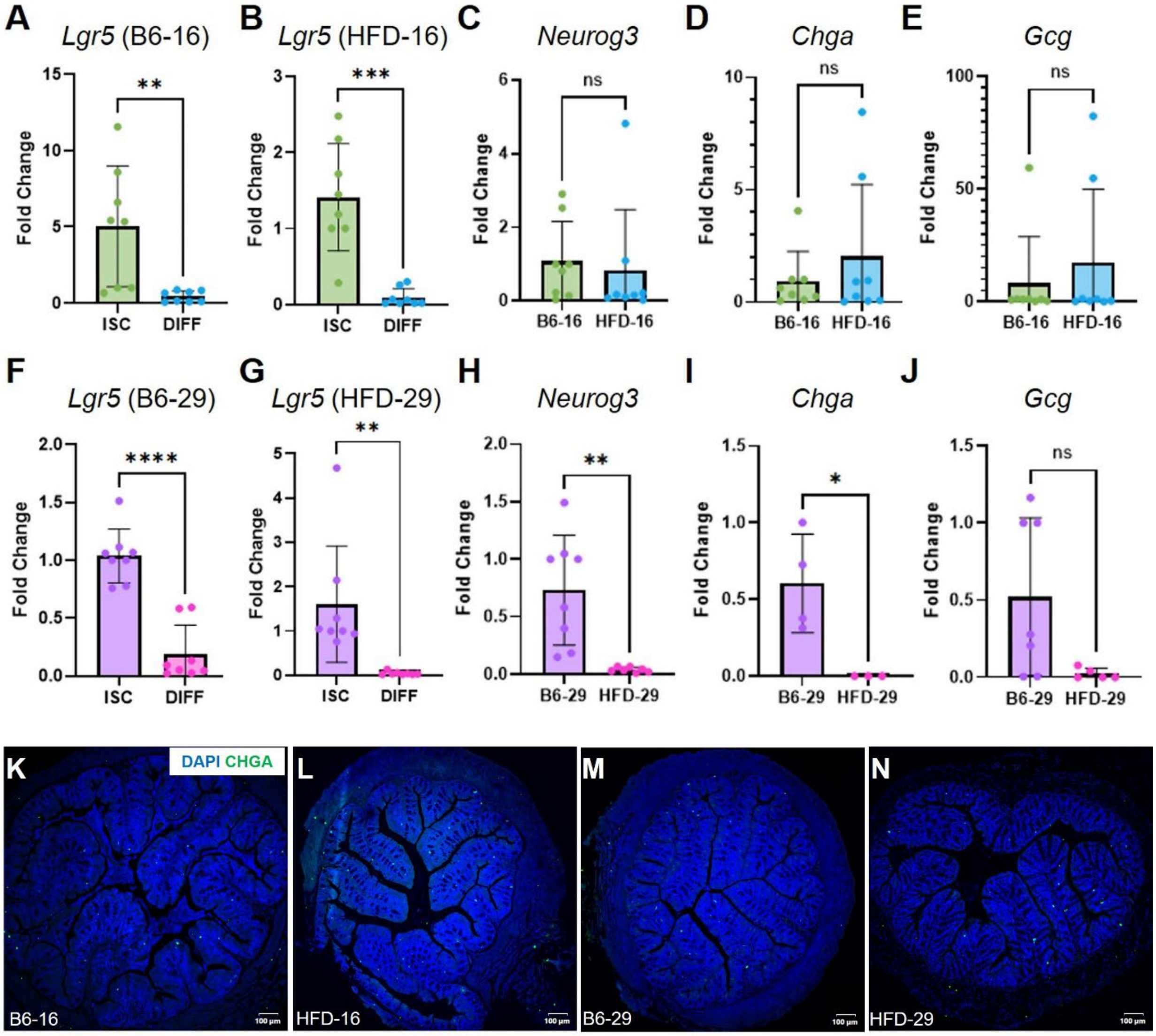


**Supplementary Figure 11. Prolonged HFD exposure impairs EEC differentiation capacity in MASH-derived colonic organoids.** Colonic crypts were isolated from control (B6-16 or B6-29) and HFD (HFD-16 or HFD-29) mice to generate colonic organoid cultures. Organoids were initially maintained in stem cell–supporting media and subsequently induced to differentiate by withdrawal of WNT3A and R-spondin. (**A**, **B**, **F**, **G**) *Lgr5* expression as detected by qPCR comparing organoids in a stem state and after differentiation. (**C**-**E**) Comparison of EEC lineage–associated genes (**C**) *Neurog3*, (**D**) *Chga*, or (**E**) *Gcg* expression between B6-16 and HFD-16 organoids. (**H**-**J**) Comparison of (**H**) *Neurog3*, (**I**) *Chga*, or (**J**) *Gcg* expression between B6-29 and HFD-29 organoids. n = 3-8 mice per group. Data analyzed using Student's t-test. *P < 0.05, **P < 0.005, ***P < 0.0005, ***P < 0.0001, NS = not significant. (**K**–**N**) Representative images of colonic cross-sections stained with DAPI (blue) and CHGA (green) from (**K**) B6-16, (**L**) HFD-16, (**M**) B6-29, (**N**) HFD-29 mice. Scale bars: 100 μm.

### Supplementary Table S1.

| Slide Number | Slide Label | Group | Diet | NASH Clinical Research Network Scoring | | | | | | |
|---|---|---|---|---|---|---|---|---|---|---|
| | | | | Macrovesicular | Microvesicular | Steatosis Grade (0-3) | Inflammation Grade (0-3) | Hepatocellular Hypertrophy (0-2) | Fibrosis Score (0-4) | Total Score |
| 1 | BL6-1-L | 16-week | Normal Chow | Negative | Negative | 0 | 0 | 0 | 0 | 0 |
| 2 | BL6-2-L | 16-week | Normal Chow | Negative | Negative | 0 | 0 | 0 | 0 | 0 |
| 3 | BL6-3-L | 16-week | Normal Chow | Negative | Negative | 0 | 0 | 0 | 0 | 0 |
| 4 | BL6-4-L | 16-week | Normal Chow | Negative | Negative | 0 | 0 | 0 | 0 | 0 |
| 5 | BL6-5-L | 16-week | Normal Chow | Negative | Negative | 0 | 0 | 0 | 0 | 0 |
| 6 | BL6-6-L | 16-week | Normal Chow | Negative | Negative | 0 | 0 | 0 | 0 | 0 |
| 7 | BL6-7-L | 16-week | Normal Chow | Negative | Negative | 0 | 0 | 0 | 0 | 0 |
| 8 | BL6-8-L | 16-week | Normal Chow | Negative | Negative | 0 | 0 | 0 | 0 | 0 |
| | | | | | | 0 | 0 | 0 | 0 | 0 |
| 11 | HFD-1-L | 16-week | High Fat Diet | Negative | (+++) | 3 | 0 | 0 | 0 | 3 |
| 12 | HFD-2-L | 16-week | High Fat Diet | (+) | (++) | 2 | 0 | 1 | 1a? | 3 |
| 13 | HFD-3-L | 16-week | High Fat Diet | (+++) | (++) | 3 | 0 | 1 | 0 | 4 |
| 14 | HFD-4-L | 16-week | High Fat Diet | (+++) | (+++) | 3 | 0 | 1 | 0 | 4 |
| 15 | HFD-5-L | 16-week | High Fat Diet | (++) | (+++) | 3 | 0 | 1 | 0 | 4 |
| 16 | HFD-6-L | 16-week | High Fat Diet | (+++) | (+++) | 3 | 0 | 1 | 0 | 4 |
| 17 | HFD-7-L | 16-week | High Fat Diet | (+) | (+++) | 3 | 0 | 1 | 0 | 4 |
| 18 | HFD-8-L | 16-week | High Fat Diet | (+) | (+++) | 2 | 0 | 0 | 0 | 2 |
| | | | | | | 2.75 | 0 | 0.75 | 0 | 3.5 |
| 21 | BL6-9-L | 29-week | Normal Chow | Negative | Negative | 0 | 0 | 0 | 0 | 0 |
| 22 | BL6-10-L | 29-week | Normal Chow | Negative | Negative | 0 | 0 (focal mild) | 0 | 0 | 0 |
| 23 | BL6-11-L | 29-week | Normal Chow | Negative | Negative | 0 | 0 | 0 | 0 | 0 |
| 24 | BL6-12-L | 29-week | Normal Chow | Negative | (+) | 1 | 0 | 1 | 0 | 1 |
| 25 | BL6-13-L | 29-week | Normal Chow | Negative | Negative | 0 | 0 | 0 | 0 | 0 |
| 26 | BL6-14-L | 29-week | Normal Chow | Negative | Negative | 0 | 0 | 0 | 0 | 0 |
| 27 | BL6-15-L | 29-week | Normal Chow | Negative | Negative | 0 | 0 | 0 | 0 | 0 |
| 28 | BL6-16-L | 29-week | Normal Chow | Negative | Negative | 0 | 0 (single focus) | 0 | 0 | 0 |
| | | | | | | 0.125 | 0 | 0.125 | 0 | 0.125 |
| 31 | HFD-9-L | 29-week | High Fat Diet | (+++) | (+) | 3 | 1 | 1 | 2 | 7 |
| 32 | HFD-10-L | 29-week | High Fat Diet | (+++) | (+) | 3 | 2 | 1 | 2 | 8 |
| 33 | HFD-11-L | 29-week | High Fat Diet | (++) | (+++) | 3 | 0 | 1 | 2 | 6 |
| 34 | HFD-12-L | 29-week | High Fat Diet | (+++) | (++) | 3 | 0 | 1 | 2 | 6 |
| 35 | HFD-13-L | 29-week | High Fat Diet | (+++) | (+++) | 3 | 0 | 1 | 0 | 4 |
| 36 | HFD-14-L | 29-week | High Fat Diet | (+++) | (+++) | 3 | 1 | 1 | 1a/1c | 6 |
| 37 | HFD-15-L | 29-week | High Fat Diet | (+++) | (+) | 3 | 0 | 1 | 0 | 4 |
| 38 | HFD-16-L | 29-week | High Fat Diet | (+++) | (+++) | 3 | 1 | 1 | 2 | 7 |
| | | | | | | 3 | 0.625 | 1 | 1.428571429 | 6 |

### Supplementary Table S2.

| Slide Number | Slide Label | Group | Diet | Submucosal Edema (0-3) | PMN Infiltration (0-3) | Goblet Cells (0-3) | Epithelial Integrity (0-3) | Total Score |
|---|---|---|---|---|---|---|---|---|
| 1 | BL6-1-C | 16-week | Normal Chow | 0 | 0 | 0 | 0 | 0 |
| 2 | BL6-2-C | 16-week | Normal Chow | 0 | 0 | 0 | 0 | 0 |
| 3 | BL6-3-C | 16-week | Normal Chow | 0 | 0 | 0 | 0 | 0 |
| 4 | BL6-4-C | 16-week | Normal Chow | 0 | 0 | 0 | 0 | 0 |
| 5 | BL6-5-C | 16-week | Normal Chow | 0 | 0 | 0 | 0 | 0 |
| 6 | BL6-6-C | 16-week | Normal Chow | 0 | 0 | 0 | 0 | 0 |
| 7 | BL6-7-C | 16-week | Normal Chow | 0 | 0 | 0 | 0 | 0 |
| 8 | BL6-8-C | 16-week | Normal Chow | 0 | 0 | 0 | 0 | 0 |
| 11 | HFD-1-C | 16-week | High Fat Diet | 0 | 0 | 0 | 0 | 0 |
| 12 | HFD-2-C | 16-week | High Fat Diet | 0 | 0 | 0 | 0 | 0 |
| 13 | HFD-3-C | 16-week | High Fat Diet | 0 | 0 | 0 | 0 | 0 |
| 14 | HFD-4-C | 16-week | High Fat Diet | 0 | 0 | 0 | 0 | 0 |
| 15 | HFD-5-C | 16-week | High Fat Diet | NA | NA | NA | NA | NA |
| 16 | HFD-6-C | 16-week | High Fat Diet | 0 | 0 | 0 | 0 | 0 |
| 17 | HFD-7-C | 16-week | High Fat Diet | 0 | 0 | 0 | 0 | 0 |
| 18 | HFD-8-C | 16-week | High Fat Diet | 0 | 0 | 0 | 0 | 0 |
| 21 | BL6-9-C | 29-week | Normal Chow | 0 | 0 | 0 | 0 | 0 |
| 22 | BL6-10-C | 29-week | Normal Chow | 0 | 0 | 0 | 0 | 0 |
| 23 | BL6-11-C | 29-week | Normal Chow | 0 | 0 | 0 | 0 | 0 |
| 24 | BL6-12-C | 29-week | Normal Chow | 0 | 0 | 0 | 0 | 0 |
| 25 | BL6-13-C | 29-week | Normal Chow | 0 | 0 | 0 | 0 | 0 |
| 26 | BL6-14-C | 29-week | Normal Chow | 0 | 0 | 0 | 0 | 0 |
| 27 | BL6-15-C | 29-week | Normal Chow | 0 | 0 | 0 | 0 | 0 |
| 28 | BL6-16-C | 29-week | Normal Chow | 0 | 0 | 0 | 0 | 0 |
| 31 | HFD-9-C | 29-week | High Fat Diet | 0 | 0 | 0 | 0 | 0 |
| 32 | HFD-10-C | 29-week | High Fat Diet | 0 | 0 | 0 | 0 | 0 |
| 33 | HFD-11-C | 29-week | High Fat Diet | 0 | 0 | 0 | 0 | 0 |
| 34 | HFD-12-C | 29-week | High Fat Diet | 0 | 0 | 0 | 0 | 0 |
| 35 | HFD-13-C | 29-week | High Fat Diet | 0 | 0 | 0 | 0 | 0 |
| 36 | HFD-14-C | 29-week | High Fat Diet | 0 | 0 | 0 | 0 | 0 |
| 37 | HFD-15-C | 29-week | High Fat Diet | 0 | 0 | 0 | 0 | 0 |
| 38 | HFD-16-C | 29-week | High Fat Diet | 0 | 0 | 0 | 0 | 0 |

### Supplementary Table S3.

| Tissue | Metabolites Significantly Upregulated | Metabolites Significantly Downregulated | Matched Alterations Between HFD-16 and HFD-29 |
|---|---|---|---|
| Liver (HFD-16 vs B6-16) | Nicotinamide, Cholic acid, Sucrose | Indole-3-ethanol, Retinal, Glycerate, Thymine, Ribothymidine, 4-Guanidinobutanoic acid, Hydroxyindoleacetic acid, N-CH3-NAM, Diaminopimelic acid, Pimelic acid, Imidazoleacetic acid, b-Aminoisobutyric acid, 4-Aminobutyrate, Histamine, Histidine, Adenosine 5’-phosphosulfate, Fumarate, Methylglutaric acid | Cholic acid (upregulated in both HFD-16 and HFD-29) |
| Liver (HFD-29 vs B6-29) | O-Phosphoethanolamine, Cholic acid, Cyclic-AMP, Glycyl-L-proline, Uracil, N-acetyl-L-asparagine | 4-Guanidinobutanoic acid, Indole-3-ethanol, Tiglic acid, Retinal, Thymine, 4-Aminobutyrate, b-Aminoisobutyric acid, 3-Nethyladipic acid, Pipecolic acid, Ribothymidine, Histidine, Hydroxyindoleacetic acid, DL-2-Amino-octanoic acid, Pimelic acid, Diaminopimelic acid | Cholic acid (upregulated in both HFD-16 and HFD-29) |
| Colon (HFD-16 vs B6-16) | Methylnicotinamide, N-Me-2PY, GTP, Propionyl-CoA, N-acetyl-L-ornithine, ATP, Uridine triphosphate, Cholic acid | Indole-3-ethanol, 4-Guanidinobutanoic acid, Hydroxyindoleacetic acid, D-Phenyllactic acid, Mannose, Diaminopimelic acid, Glycyl-L-leucine, Hydroquinone, N-acetyllysine, Dimethylmalonic acid | Cholic acid (upregulated in both HFD-16 and HFD-29) |
| Colon (HFD-29 vs B6-29) | N-acetyl-L-proline, Glycerate, Cholic acid, Thymine, Imidazoleacetic acid hydrochloride, N-acetyl-L-ornithine | 4-Guanidinobutanoic acid, D-Phenyllactic acid, Glutaric acid, Hydroquinone, Dimethylmalonic acid, Hydroxyindoleacetic acid, 3-Hydroxybutyrate, Indole-3-ethanol, Butyric acid, Lactose, 3-Methyladipic acid, NADPH, N-acetyllysine, Glycyl-L-leucine, DL-2-Amino-octanoic acid, Uridine triphosphate | Cholic acid (upregulated in both HFD-16 and HFD-29) |
| Stool (HFD-16 vs B6-16) | 2-PY, Cytidine, IMP, 6-Hydroxymelatonin glucuronide, Threonine, Carnitine, Cholic acid, Acetoacetate, Creatine, Aspartate | Kynurenic acid, Melatonin, Hydroxyindoleacetic acid, Glutathione disulfide, N-Octanoylglycine, 4-Pyridoxic acid, Kynurenine, Ribose, N-acetyl-L-isoleucine, DL-2-Aminooctanoic acid, Hydroquinone, N-acetyltyrosine, N-CH3-NAM, N-acetylserotonin, Serotonin, Lactose, Glucose, Pimelic acid, dUMP, Malonic acid, Cellobiose, Tryptamine, Sucrose, Malate, Suberic acid, N-acetyl-L-leucine, Galacturonic acid, D-gluconate, Sepiapterin, Guanadrel sulfate, 5-Methylthioadenosine, Quinolinate, Hydroxyphenyllactic acid, L-2-Hydroxyglutaric acid, Deoxyadenosine, Methylnicotinamide, D-Phenyllactic acid, D-erythrose-4-phosphate, N-acetyl-L-glutamine, Imidazoleacetic acid | Cholic acid (upregulated in both HFD-16 and HFD-29) |
| Stool (HFD-29 vs B6-29) | Cholic acid, IMP, Cytosine, Biotin, 2-PY, Anthranilate, Acetyl-L-asparagine, ADP-D-glucose, Leucine, Thymine, N-Me-2PY, Phosphoserine, Aspartate, Guanidoacetic acid | Melatonin, N-acetylserotonin, N-Octanoylglycine, Hydroxyindoleacetic acid, 4-Pyridoxic acid, Guanadrel sulfate, N-acetyl-L-isoleucine, Tryptamine, 5-Methoxytryptamine, Hydroquinone, N-acetyltyrosine, NADH, 5-Methoxytryptophan, D-gluconate, Kynurenic acid, Glucose, Glutaric acid, 5’-Methylthioadenosine, N-acetyl-L-leucine, Glutathione disulfide, Sepiapterin, Pimelic acid, Ribose, N-CH3-NAM, D-Phenyllactic acid, Lactate, Kynurenine, Lactose, 2-Furoylglycine, Hydroxyphenyllactic acid, Butyric acid, Tiglic acid, Deoxyadenosine, Malate, dUMP, Taurodeoxycholic acid, Ribose-phosphate, Pyridoxine, Hydroxyphenylpyruvate, Methylnicotinamide, Aminolevulinic acid | Cholic acid (upregulated in both HFD-16 and HFD-29) |

### Supplementary Table S4.

| Abbreviation | Common Name | Main Class | Lipid Category |
|---|---|---|---|
| ADGGA | Acyl diacylglyceryl glucuronide | Other glycerolipids [GL00] | Glycerolipids (GL) |
| AHexCer | Acylhexosylceramide | Neutral glycosphingolipids [SP05] | Sphingolipids (SP) |
| ASG | Acylsterylglycosides | Other Sterol Lipids | Sterol Lipids (ST) |
| BA | Cholic Acid | Bile acids and derivatives [ST04] | Sterol Lipids (ST) |
| CAR | Acylcarnitine | Fatty esters [FA07] | Fatty acyls (FA) |
| CE | Cholesteryl esters | Sterols [ST01] | Sterol Lipids (ST) |
| Cer | Ceramide | Ceramides [SP02] | Sphingolipids (SP) |
| CerP | Ceramide-1-phosphate | Ceramides [SP02] | Sphingolipids (SP) |
| CL | Cardiolipins | Glycerophosphoglycerophosphoglycerol [GP12] | Glycerophospholipids (GP) |
| DG | Diacylglycerol | Diradylglycerols [GL02] | Glycerolipids (GL) |
| DGGA | Diacylglyceryl glucuronide | Other glycerolipids [GL00] | Glycerolipids (GL) |
| DLCL | Dilysocardiolipin | Glycerophosphoglycerophosphoglycerol [GP12] | Glycerophospholipids (GP) |
| FA | Free fatty acid | Fatty acids and Conjugates [FA01] | Fatty acyls (FA) |
| FAHFA | Fatty acid ester of hydroxyl fatty acid | Fatty esters [FA07] | Fatty acyls (FA) |
| GM3 | Ganglioside | Acidic glycosphingolipids [SP06] | Sphingolipids (SP) |
| HBMP | Hemibismonoacylglycerophosphate | Glycerophosphoglycerols [GP04] | Glycerophospholipids (GP) |
| Hex2Cer | Dihexosylceramide | Neutral glycosphingolipids [SP05] | Sphingolipids (SP) |
| Hex3Cer | Trihexosylceramide | Neutral glycosphingolipids [SP05] | Sphingolipids (SP) |
| HexCer | Hexosylceramide | Neutral glycosphingolipids [SP05] | Sphingolipids (SP) |
| LPA | Lysophosphatidic acid | Glycerophosphates [GP10] | Glycerophospholipids (GP) |
| LPC | Lysophosphatidylcholine | Glycerophosphocholines [GP01] | Glycerophospholipids (GP) |
| LPE | Lysophosphatidylethanolamine | Glycerophosphoethanolamines [GP02] | Glycerophospholipids (GP) |
| LPE-N | N-acyl-lysophosphatidylethanolamine (lyso-NAPE) | Glycerophosphoethanolamines [GP02] | Glycerophospholipids (GP) |
| LPG | Lysophosphatidylglycerol | Glycerophosphoglycerols [GP04] | Glycerophospholipids (GP) |
| LPI | Lysophosphatidylinositol | Glycerophosphoinositols [GP06] | Glycerophospholipids (GP) |
| LPS | Lysophosphatidylserine | Glycerophosphoserines [GP03] | Glycerophospholipids (GP) |
| MG | Monoacylglycerol | Monoradylglycerols [GL01] | Glycerolipids (GL) |
| MGDG | Monogalactosyldiacylglycerol | Glycosyldiradylglycerols [GL05] | Glycerolipids (GL) |
| MLCL | Lysocardiolipin | Glycerophosphoglycerophosphoglycerol [GP12] | Glycerophospholipids (GP) |
| NAE | N-acyl ethanolamines | Fatty amides [FA08] | Fatty acyls (FA) |
| NAGly | N-acyl glycine | Fatty amides [FA08] | Fatty acyls (FA) |
| NAGlySer | N-acyl glycyl serine | Fatty amides [FA08] | Fatty acyls (FA) |
| NATau | N-arachidonoyl taurine | Fatty amides [FA08] | Fatty acyls (FA) |
| PA | Phosphatidic acid | Glycerophosphates [GP10] | Glycerophospholipids (GP) |
| PC | Phosphatidylcholine | Glycerophosphocholines [GP01] | Glycerophospholipids (GP) |
| PE | Phosphatidylethanolamine | Glycerophosphoethanolamines [GP02] | Glycerophospholipids (GP) |
| PE-Cer | Ceramide phosphoethanolamine | Phosphosphingolipids [SP03] | Sphingolipids (SP) |
| PEtOH | Phosphatidylethanol | Glycerophospholipids [GP10] | Glycerophospholipids (GP) |
| PG | Phosphatidylglycerol | Glycerophosphoglycerols [GP04] | Glycerophospholipids (GP) |
| PI | Phosphatidylinositol | Glycerophosphoinositols [GP06] | Glycerophospholipids (GP) |
| PI-Cer | Ceramide phosphoinositol | Phosphosphingolipids [SP03] | Sphingolipids (SP) |
| PMeOH | Phosphatidylmethanol | Other Glycerophospholipids [GP00] | Glycerophospholipids (GP) |
| PS | Phosphatidylserine | Glycerophosphoserines [GP03] | Glycerophospholipids (GP) |
| PT | Diacylglycerophosphothreonines | Glycerophosphothreonines [GP24] | Glycerophospholipids (GP) |
| SE | Steryl ester | Sterols [ST01] | Sterol Lipids (ST) |
| SHexCer | Stigmasterol hexoside | Sterols [ST01] | Sterol Lipids (ST) |
| SL | Sulfonolipid | Sphingoid bases [SP01] | Sphingolipids (SP) |
| SM | Sphingomyelin | Phosphosphingolipids [SP03] | Sphingolipids (SP) |
| SMGDG | Semino Lipid | Glycosyldiradylglycerols [GL05] | Glycerolipids (GL) |
| SPB | Sphingoid base | Sphingoid bases [SP01] | Sphingolipids (SP) |
| SQDG | Sulfoquinovosyl diacylglycerol | Glycosyldiradylglycerols [GL05] | Glycerolipids (GL) |
| ST | Sterol | Sterols [ST01] | Sterol Lipids (ST) |
| TG | Triacylglycerol | Triradylglycerols [GL03] | Glycerolipids (GL) |
| VAE | Vitamin A fatty acid ester | Isoprenoids [PR01] | Prenol lipids (PR) |

### Supplementary Table S5.

| 1° Antibodies | Host | Company | Ref # | Lot # |
|---|---|---|---|---|
| CHGA | Rabbit / IgG | abcam | ab45179 | 1060116-1 |
| 5-HT | Goat | abcam | ab66047 | 1046050-21 |
| Fibronectin | Mouse | Invitrogen | MA5-11981 | ZA4197162 |

| 2° Antibodies | Host | Company | REF | Lot # |
|---|---|---|---|---|
| Donkey anti Goat 488 | Donkey | Invitrogen | A11055 | 1869589 |
| Goat anti Rabbit 594 | Goat | Invitrogen | A11012 | 2616076 |
| Donkey anti Mouse 488 | Donkey | Southern Biotech | 6411-30 | L1013-NK00Z |
| Goat anti Mouse 561 | Goat | Invitrogen | A11005 | 2641993 |
| DAPI | | Invitrogen | 62248 | YD3918381 |

### Supplementary Table S6.

| Company | Gene Name | Lot # | Catalog # |
|---|---|---|---|
| ThermoFisher | Gcg | 2160159 | 4448892 |
| ThermoFisher | Neurog3 | 1693792 | 4331182 |
| ThermoFisher | Chga | P241011-005 H09 | 4331182 |
| ThermoFisher | Actb | P240724-012 F11 | 4331182 |
| ThermoFisher | Lgr5 | P240417-006 H03 | 4448892 |
